# Regulatory networks driving expression of genes critical for glioblastoma are controlled by the transcription factor c-Jun and the pre-existing epigenetic modifications

**DOI:** 10.1101/2022.07.18.500476

**Authors:** Adria-Jaume Roura, Paulina Szadkowska, Katarzyna Poleszak, Michal J. Dabrowski, Aleksandra Ellert-Miklaszewska, Kamil Wojnicki, Iwona A. Ciechomska, Karolina Stepniak, Bozena Kaminska, Bartosz Wojtas

## Abstract

**Background:** Glioblastoma (GBM, WHO grade IV) is an aggressive, primary brain tumor. Despite gross surgery and forceful radio- and chemotherapy, survival of GBM patients did not improve over decades. Several studies reported transcription deregulation in GBMs but regulatory mechanisms driving overexpression of GBM-specific genes remain largely unknown. Transcription in open chromatin regions is directed by transcription factors (TFs) that bind to specific motifs, recruit co-activators/repressors and the transcriptional machinery. Identification of GBM-related TFs-gene regulatory networks may reveal new and targetable mechanisms of gliomagenesis.

**Results:** We predicted TFs-regulated networks in GBMs *in silico* and intersected them with putative TF binding sites identified in the accessible chromatin in human glioma cells and GBM patient samples. The Cancer Genome Atlas and Glioma Atlas datasets (DNA methylation, H3K27 acetylation, transcriptomic profiles) were explored to elucidate TFs-gene regulatory networks and effects of the epigenetic background. In contrast to the majority of tumors, c-Jun expression was higher in GBMs than in normal brain and c-Jun binding sites were found in multiple genes overexpressed in GBMs such as *VIM, FOSL2* or *UPP1*. Binding of c-Jun to the *VIM* gene promoter is stronger in GBM cells than in cells derived from benign glioma as evidenced by gel shift and supershift assays. Regulatory regions of a majority of the c-Jun targets have distinct DNA methylation in GBMs suggesting the contribution of DNA methylation to the c-Jun-dependent regulation.

**Conclusions:** We identified distinct TFs-gene networks in GBMs compared to benign gliomas, a predominant role of c-Jun in controlling genes driving gliomagenesis and a modulatory role of DNA methylation.

## Background

Malignant gliomas represent 80% of malignant tumors of the central nervous system (CNS) and are most common primary CNS tumors in adults. An integrated classification introduced by the World Health Organization (WHO) in 2016 categorized gliomas into malignancy grades (I-IV) based on tumor morphology and molecular information [1,2]. Comprehensive transcriptomic, genomic and epigenetic analyses showed gross differences between low-grade gliomas (LGGs, WHO grade I-II) and high-grade gliomas (HGGs, WHO grade III-IV) [2]. Glioblastoma (GBM) is a highly malignant, diffuse tumor with a poor prognosis (mean patient survival is 15 months) due to a lack of efficacy of both conventional and emerging therapies, including immune checkpoint blockade therapies. Deregulation of transcription due to aberrant activation of signaling pathways is manifested by activation, repression and/or temporal/spatial deregulation of many genes [3], all of which contributes to tumor initiation and progression. We have recently demonstrated the widespread deregulation of chromatin accessibility, histone modifications and transcriptional profiles in GBMs in comparison to benign gliomas [4].

A crucial step in gene expression regulation is initiation of transcription, which is strictly regulated by DNA-binding proteins known as transcription factors (TFs) which bind to gene promoters and enhancers [5]. Altered TF activities had been linked to a variety of cancers, with an estimated 20% of oncogenes being identified as TFs [6]. TFs bind mainly in open chromatin regions and usually cooperate with other DNA-binding proteins to regulate gene expression [7,8]. Different mechanisms such as gene amplifications, point mutations, expression changes along with DNA methylation or histone modifications can influence TF activities in cancer [9]. Overexpression of certain TFs may serve as a prognostic marker in malignant gliomas [10].

Initial ENCODE studies revealed three main TFs localizing almost exclusively within accessible chromatin: c-Jun, GATA1 and NRF1 [11]. c-Jun is a component of the Activator protein-1 (AP-1), a dimer composed of proteins of the Jun (c-Jun, JunB, JunD), and Fos (c-Fos, FosB, FRA-1 and FRA-2) families [12]. The AP-1 complex comprising c-Jun stimulates cell proliferation through the repression of tumor suppressor genes [13] or the induction of *CYCLIN D1* [14–16]. JunB and JunD act frequently as negative regulators [17,18].

To identify TF-gene regulatory networks driving transcriptional deregulation in gliomas, we exploited public TCGA datasets as well as the Glioma Atlas created in our laboratory, which encompasses genome-wide profiles of chromatin accessibility, histone modifications, DNA methylation and gene expression from the same tumor sample [4] (presented as a summary at the Additional file: Fig. S1). Intersecting the acquired data resulted in mapping of active promoters and enhancers in gliomas, and identification of GBM-specific active sites [4]. We combined these multiple datasets with data on chromatin accessibility, H3K27 acetylation and gene expression in two cultured human glioblastoma cell lines to predict gene regulatory networks and select candidate TFs enriched in GBMs. We discovered a set of TFs, including c-Jun, that likely control genes implicated in tumor progression in GBMs. We verified the binding and presence of c-Jun at two candidate gene promoters (*VIM, UPP1)* in glioma cells and found a stronger binding in glioblastoma cells in comparison to cells derived from the low-grade glioma. We demonstrate that GBM-specific DNA methylation patterns exist in c-Jun target genes and verify that methylation in the c-Jun binding site may affect its binding. Our findings show how specific TF-gene regulatory networks contribute to malignant glioma pathogenesis.

## Results

### Identification of TF binding sites in open chromatin regions specific to malignant gliomas

First, we predicted transcription factor binding sites (TFBS) from chromatin accessibility peaks (ATAC-seq) identified in LN18 and LN229 human glioma cells. Only peaks consistently detected in both cell lines were considered. Secondly, we compared the obtained results with the ATAC-seq data from GBMs (n=2) [4]. Data were acquired from a genome browser [19]. At least 60% overlapping TFBS calls were identified between samples (Fig. 1A). The TFBS detected in both cell lines (145,123) or GBMs (219,352) were further analyzed. Chromatin accessible regions were noticeably different in cultured glioma cells and tumor samples (Fig. 1A), which may reflect the cellular heterogeneity of GBMs, composed of tumor cells, stromal cells and numerous infiltrating immune cells (up to 40%). We created heatmaps of all detected peaks to identify the ATAC-seq signal enrichment near the transcription start site (TSS), and evaluated the precise distribution of peaks in these promoters. Open chromatin peaks occurred mostly in the vicinity of the TSS (Fig. 1B).

**Fig. 1.**
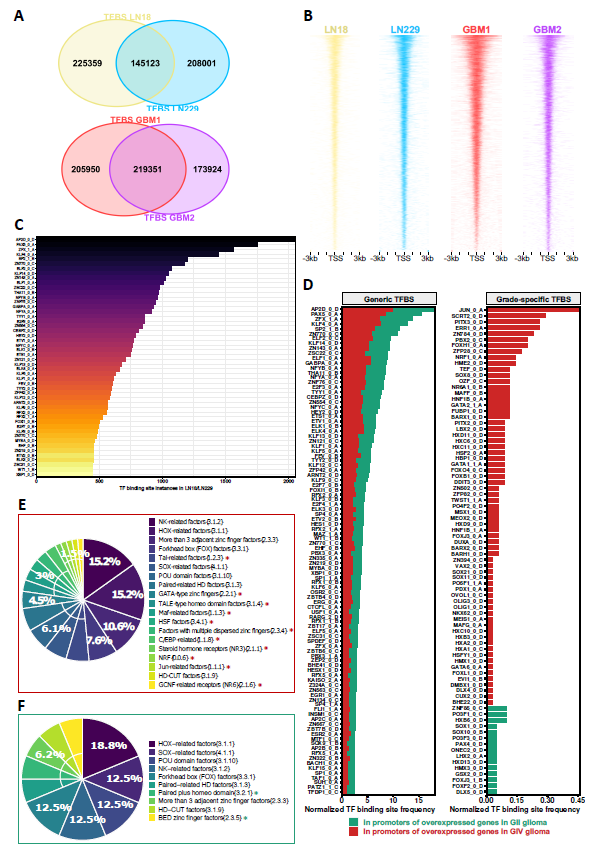
Global characterization of transcription factor binding sites in open chromatin regions in glioblastoma cell lines and glioblastoma specimens. **(A)** Total number of predicted transcription factor binding sites in open-chromatin regions using ATAC-seq fragments and position-weight matrices (PWMs) motifs in established human glioma cell lines LN18 and LN229 and in glioblastoma samples. In-silico bound predictions for both cell lines were selected for downstream analysis. **(B)** Profile heatmap of total ATAC-seq peaks identified around transcription start sites (TSS) in cell-lines and GBM specimens. **(C)** Top 50 occurrences of transcription factor binding sites (HOCOMOCOv11) in gene promoters (TSS±1.5kb) in LN18 and LN229 glioma cell lines; the last letter (A-D) represents the quality, where A represents motifs with the highest confidence and the number defines the motif rank, with zero indicating the primary model (primary binding preferences). **(D)** Prediction of “generic” (left-panel) and “grade-specific” (right-panel) transcription factor binding sites in the promoters (TSS±1.5kb) of dysregulated genes in gliomas of grade IV and II. The abscissa represents a normalization factor for TFBS occurrences in which the total number of differentially expressed genes in a given glioma grade is taken into account. **(E-F)** Transcription factor (TF) families (HOCOMOCOv11) with putative binding sites in the promoters of overexpressed genes in IV glioma **(E)** and grade II glioma **(F)**. Unique TF families found in either group are highlighted with asterisks.

In glioma cell lines, we found that TFBS in open-chromatin regions of the genome contain numerous motifs for the AP2D, PAX5, and ZFX binding proteins, among many others (Fig. 1C). We performed a differentially expressed gene (DEG) analysis using TCGA data (Additional file 2: Fig. S1) to identify genes that were either overexpressed in GIV gliomas or in benign gliomas (WHO grade II gliomas, GII). Next, we searched for TFBS only within the promoter regions of these genes (TSS±1.5 kb). We found 3,454 genes overexpressed in GBMs and 2,010 genes overexpressed in GII gliomas (Additional file 2: Fig. S1). We found TFBS in the promoter regions of these genes and labelled them “generic TFBS” if they were found in any set of overexpressed genes, and “grade-specific TFBS” if they were only identified in one set (Fig. 1D). “Generic TFBS” motifs were in equal proportions in the GIV and GII specific gene promoters, indicating that these motifs could be engaged in housekeeping or brain specific functions (Fig. 1D). However, we found 240 TFBS that were only found in the promoters of genes overexpressed in glioblastomas (GIV), and this set included binding sites for c-Jun, SCRT2, PITX3, ERR1, or ZN784. TFBS for factors such as ZNF85, PO3F1, HX36, SOX1, and SOX10 were found in genes overexpressed in GII gliomas (Fig. 1D).

We annotated these grade-specific transcription factors into TF families using the HOmo sapiens COmprehensive MOdel COllection (HOCOMOCO) v11 database [20], and found that specific TF protein families may be more relevant for transcription regulation in GIV or GII gliomas (Fig. 1E, F). Binding sites for TFs belonging to families such as Tal-related factors, GATA-type zinc fingers, or Jun-related factors were present only in the genes overexpressed in GIV gliomas (Fig. 1E). Motifs for some TF families, such as NK-related factors, HOX-related factors, and POU-related factors, were enriched both in GII and GIV gliomas. These findings suggest that TFs from particular TF families drive gene expression networks deregulated in GIV gliomas.

Next, we examined transcriptomic differences between GIV and GII gliomas from the TCGA dataset (Additional file 2: Fig. S2A). Patient samples clustered depending on a glioma grade (Additional file 2: Fig. S2A) in agreement with previous studies [21–23]. Numerous transcriptomic differences between GBMs and LGGs were detected (Additional file 2: Fig. S2B). The genes overexpressed in LGGs were functionally related to synaptic and neuronal functions (Additional file 2: Fig. S2C-D), whereas in GBMs overexpressed genes were related to immune responses and cell cycle, as shown by the Gene Set Enrichment Analysis (GSEA). Furthermore, a pathway enrichment analysis revealed that genes up-regulated in GBMs were associated with the p53 signaling pathway, cell cycle, IL-17 signaling, nucleosome assembly, and extracellular matrix (ECM) organization, whereas genes up-regulated in LGGs were associated with neuroactive interactions, GABAergic synapses and synaptic plasticity (Additional file 2: Fig. S2E-F).

### *c-Jun* expression in malignant gliomas is deregulated in an opposite way than in most cancers

The above presented findings show that c-Jun transcription factor is predicted to bind to the promoters of the genes overexpressed in GBMs (Fig. 1D). We used the Pan-Cancer and glioma TCGA datasets to investigate *JUN* expression in various types of cancers and corresponding normal tissues (Fig. 2A). In most cancers (i.e., BLCA, BRCA, SKCM, CESC, OV, LUSC, UCEC, LUAD, and UCS; full name in Methods), *JUN* expression was significantly lower in tumors than in non-tumor tissues. Only in two tumor types, i.e., thymoma (THYM) and GBM *JUN* expression was increased as compared to neighboring tissues. Moreover, *JUN* expression increased along with the malignant progression and was the highest in GBMs in the TCGA dataset (Fig. 2B).

**Fig. 2.**
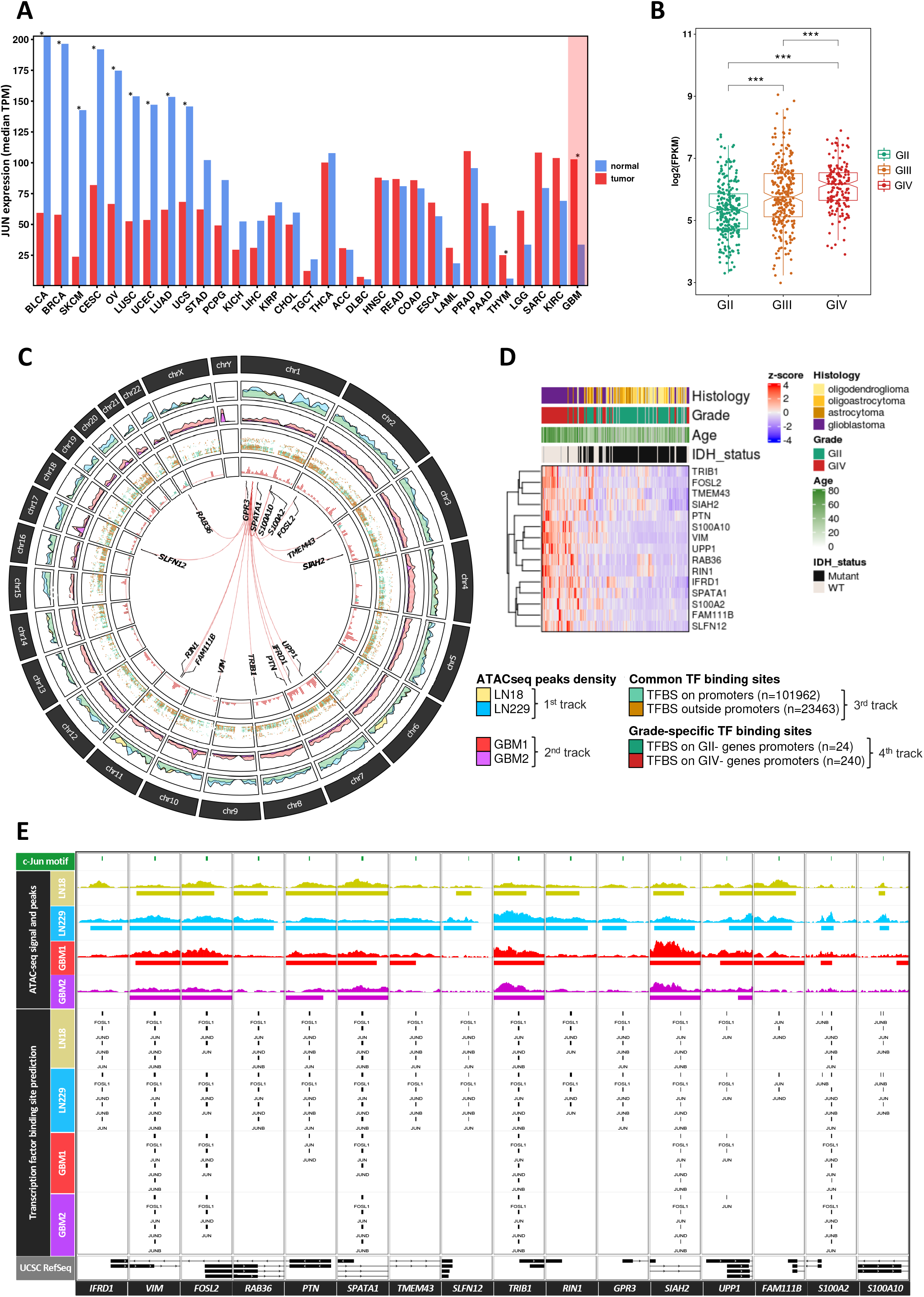
c-Jun dysregulation in the Pan-cancer atlas and the TCGA glioma dataset and identification of genomic targets in open-chromatin regions. **(A)** JUN mRNA expression profile ordered by expression differences between tumor samples (TCGA) and paired normal tissues (TCGA normal + GTEx normal). The differential expression was calculated using one-way ANOVA and disease state (Tumors or Normal, *p-value < 0.05). **(B)** c-Jun mRNA expression across glioma grades using the TCGA data. The differential expression was calculated using Wilcoxon rank-sum statistical test (*p-value < 0.05, **p-value < 0.01 and ***p-value < 0.001). **(C)** Chromatin accessibility profiling (1^st^ and 2^nd^ tracks) and TFBS predictions in or outside promoters (3^rd^ track). The 4th track depicts TFBS predictions on over-expressed genes in certain glioma grades, and red lines connect JUN’s gene location (*chr1:58776845-58784048*) to each of the c-Jun-controlled genes in GBM. **(D)** Unsupervised hierarchical clustering heatmap of c-Jun gene targets in grade II and grade IV gliomas from the TCGA dataset (248 grade II gliomas; 160 grade IV gliomas). **(E)** Landscape of c-Jun binding prediction in the cis-regulatory regions of selected over-expressed GIV genes, in the studied cell lines and GBM patient samples. Location of the identified c-Jun motif is shown in green. The ATAC-seq signal and MACS2 broad peaks for each cell line and GBM patient sample are shown separately. Exons (rectangles) and introns (lines) are depicted as well as the gene orientation (arrows) in the UCSC gene composite track.

### Identification of c-Jun-gene regulatory networks in cultured human glioma cells

Patterns of ATAC-seq peaks, representing chromatin accessible regions, were consistent in LN18 and LN229 glioma cells (Fig. 2C, 1st track) and showed a considerable similarity to patterns of accessible chromatin identified in GBMs (Fig. 2C, 2nd track). After intersecting ATAC-seq peaks with the H3K27 acetylation peaks, we identified 101,962 TFBS in promoter regions (TSS±1.5 kb), accounting for ∼81.3% of all TFBS predictions; whereas only ∼18.7% of TFBS were found outside of the promoter regions (Fig. 2C, 3rd track).

In the promoters of genes overexpressed in GII gliomas (Additional file 2: Fig. S1), we detected 24 putative TFBS, while in the promoters of genes overexpressed in GBMs, we found 240 putative TFBS (Fig. 2C, 4th track). Interestingly, many c-Jun binding sites were found in the promoters of genes involved in immune-related signaling (*IFRD1, UPP1*, and *SLNF12)*, cell proliferation, migration, invasion (*VIM, FOSL2, PTN, SIAH2, S100A2, S100A10* and *FAM111B)* and radio-resistance (*TRIB1)*. All of these genes were significantly up-regulated in GBMs when compared with GII (Fig. 2D). Several Jun-related factors and Fos-related factors can bind to the same binding sites as c-Jun within regulatory regions of potential c-Jun targets (Fig. 2E). Similar putative TFBS were identified in glioma cells and GBMs in 50% (8/16) of the gene promoters (Fig. 2E, GBM1 and GBM2 TFBS predictions).

We studied if patient survival is associated with the expression of *JUN* and its target genes in GBM-TCGA (Additional file 2: Fig. S3) and LGG-TCGA samples (Additional file 2: Fig. S4). High expression of c-Jun targets: *FOSL2, GPR3, RIN1* and *UPP1* (Log-rank p-values<0.05) was associated with a worse prognosis. Patient survival analysis of the LGG patients showed that high expression of c-Jun targets was associated with the worse prognosis.

### Transcriptomic analysis of grade-specific transcription factors

The enrichment of TFBS in open-chromatin regions in GIV gliomas (red bars on the right panel in Fig. 1D) suggested that the corresponding TFs regulate genes important for glioma progression. We examined expression of GBM specific TF encoding genes (64 genes) using hierarchical clustering of TCGA GII and GIV (Fig. 3A). We found that while most of them are highly expressed in GIV, some are more prominent in GII. Many HOMEOBOX (HOX)-related genes (*HOXD11, HOXD9, HOXC10, HOXC11, HOXC6, HOXB3, HOXA2, HOXA1*) were associated with a glioma grade and significantly overexpressed in GBMs (Wilcoxon rank-sum and BH padj <0.05). *HOX* genes are involved in developmental processes [24] so they are not expressed in adult brain, but they are re-expressed in malignant gliomas and linked to tumorigenesis [25]. To produce gene-specific regulatory outcomes, interactions of HOX factors with other TFs (that are expressed in tissue- and cell type-specific manners) are required [26]. Expression of *JUN* was significantly up-regulated in GIV when compared to GII (Fig. 3A). Genes coding for TFs upregulated in GBMs (Fig. 3B) were statistically significantly overexpressed in 90% high-grade gliomas (27/30), whereas genes coding for TFs that were associated with GII (Fig. 3B) were significantly overexpressed across glioma grades in only 53.15% (17/32) cases. Factors such as *MEOX2, TWIST1, MAFF, DDIT3, MEIS1* were overexpressed in GIV. A number of TF coding genes was specifically overexpressed in GII (Fig. 3B).

**Fig. 3.**
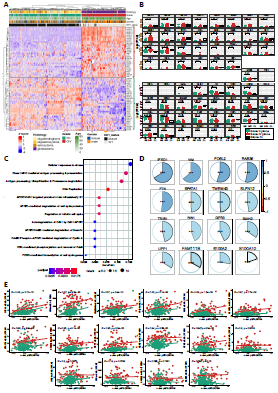
Transcription factor expression differs across glioma grades and c-Jun positively correlates with its target genes. **(A)** Unsupervised clustering of genes coding for grade-specific transcription factors. The TCGA patients (grade II: 248 patients, grade IV: 160 patients) and genes were clustered using Ward’s minimum variance method. Patients who lack clinical information on Histology, Grade, Age or Gender are illustrated in grey. **(B)** Normalized transcription factor expression in grade II and grade IV glioma (TCGA RNA-seq data) and in established glioma cell lines LN18 and LN229 (CL; 2 replicates of each shown). TFs were grouped based on dendrogram clusters depicted in A. The adjusted p-values for statistical differences between glioma grades are displayed and the Wilcoxon rank-sum statistical test and Benjamini-Hochberg (BH) correction were used. Transcription factors with a logarithmic expression of zero or nearly zero in glioma patients have no statistical validity. **(C)** Reactome analysis of genes having grade IV-specific transcription factor motifs in their promoters. BH procedure was used to correct for multiple testing. **(D)** Correlation of mRNA levels between JUN mRNA and its targets (TCGA grade II and grade IV patients). Genes are ordered based on obtained Pearson’s correlation (blue to red scale) and associated p-values were corrected by multiple testing (*padj < 0.05, **padj < 0.01 and ***padj < 0.001). **(E)** Pearson’s correlation coefficient of c-Jun reverse phase protein array (phosphorylated c-Jun pS73) against mRNA of c-Jun target genes. The adjusted BH p-values for statistically significant correlations are displayed. The data points are color-coded according to the glioma’s grade, along with their regression line.

The expression of preselected TFs was also evaluated in RNA-seq data from human LN18 and LN229 glioma cells (Fig. 3B, black boxplots). Gene expression medians for many genes encoding TFs were consistent with the patterns detected in the tumor samples (Fig. 3B). Genes coding for transcription factors PDX1, OLIG3, POU5F1, PITX3, FOXH1, OVOL1, GATA1, HNF1B, BARX2, POU42F2 were expressed at a very low level (Fig. 3B). Even though their motifs were found in the promoters of GIV-related genes, expression of associated TFs might be lost in cultured cells or they have specific expression in non-tumor cells in GBMs.

Focusing on GIV-specific TFs (Fig. 1D, right panel), we selected 166 genes that have at least one TF motif predicted in the gene promoter and were significantly upregulated in GIV compared to GII (student’s t-test and FDR<0.05). Then, to better understand biological functions of these genes, we performed a pathway enrichment analysis (Fig. 3C). We found that many of these genes are involved in cellular stress responses, DNA replication, cell cycle, and antigen processing and presentation. This suggests that the identified GIV-specific TFs may influence expression of critical genes involved in glioma progression.

### Expression of *JUN* positively correlates with expression of putative targets

We hypothesized that high levels of c-Jun will result in the increased expression of its target genes. We calculated the correlation between the *JUN* mRNA level and the expression of c-Jun targets in LGGs and GBMs in the TCGA dataset (Fig. 3D). We found a positive and significant Pearson’s correlation (adjusted p-values<0.05) in all of the cases, with the highest positive correlation for genes encoding interferon related developmental regulator 1 (*IFRD1*), VIMENTIN (*VIM*), and FOS Like 2 (*FOSL2*). Using publicly available reverse protein phase assay (RPPA) data [27], we compared the level of the phosphorylated c-Jun (serine 73, S73) and expression of sixteen genes in the glioma samples (Fig. 3E). The higher levels of phosphorylated c-Jun significantly correlated with mRNA levels of putative c-Jun targets. Based on the mRNA-to-mRNA correlation (Fig. 3D) as well as the phosphorylated c-Jun-to-mRNA correlation, *FOSL2* and *VIM* were the most positively correlated targets (Fig. 3E).

### Motifs for c-Jun and other basic leucine zipper proteins are enriched in glioma distal-regulatory regions

We had previously identified enhancers enriched in active histone H3K27ac marks in topologically associating domains (TADs) in GI (pilocytic astrocytomas, PAs), and HGGs (diffuse astrocytomas, DAs and GBMs) [4]. In the present analysis, we focused on common active enhancers found in several glioma patients (Fig. 4A, 1st track). Subsequently, we intersected all predicted TF motifs within these regulatory regions, which yielded 7,571 TF motif instances (Fig. 4A, 2nd track). A total of 94 binding sites for the c-Jun were identified within glioma enhancers (Additional file 1: Table S1).

**Fig. 4.**
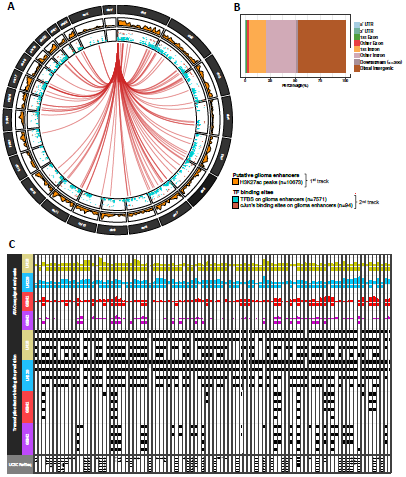
The integration of the glioma enhancer atlas in the context of c-Jun and related factors. **(A)** The density of the ChIPseq H3K27ac peak from the putative glioma enhancer atlas (1^st^ track) identified by Stepniak et al. TFBS motif predictions in LN18 and LN229 glioma cell lines inside enhancers and c-Jun binding sites are displayed separately (2^nd^ track). Each putative JUN motif found in glioma enhancers is linked to *JUN* gene position (*chr1:58776845-58784048*). **(B)** Feature distribution of glioma enhancer (putative H3K27ac peaks). **(C)** Integration of glioma enhancers and chromatin openness in glioma cell lines and GBM specimens with TFBS for c-Jun and other bZIP proteins. Exons (rectangles) and introns (lines) are depicted as well as the gene orientation (arrows) in the UCSC gene composite track.

We evaluated cumulative hypergeometric probabilities to quantify the enrichment of particular TFBS within glioma enhancers and discovered that several basic leucine zipper (bZIP) transcription factors, including c-Jun, are found significantly at the top of our TFBS ranking (Table 1, Additional file 1: Table S2). This finding suggests that in gliomas, the bZIP TF class, which comprises the Fos-, Jun- and Maf-related families, is involved in gene regulation via promoters and enhancers. Indeed, consensus H3K27ac peaks in GBMs were primarily observed in distal intergenic regions, followed by intronic regions (Fig. 4B). This finding shows that many intron DNA sequences may contain important elements for aberrant transcription control in tumors.

**Table 1:**
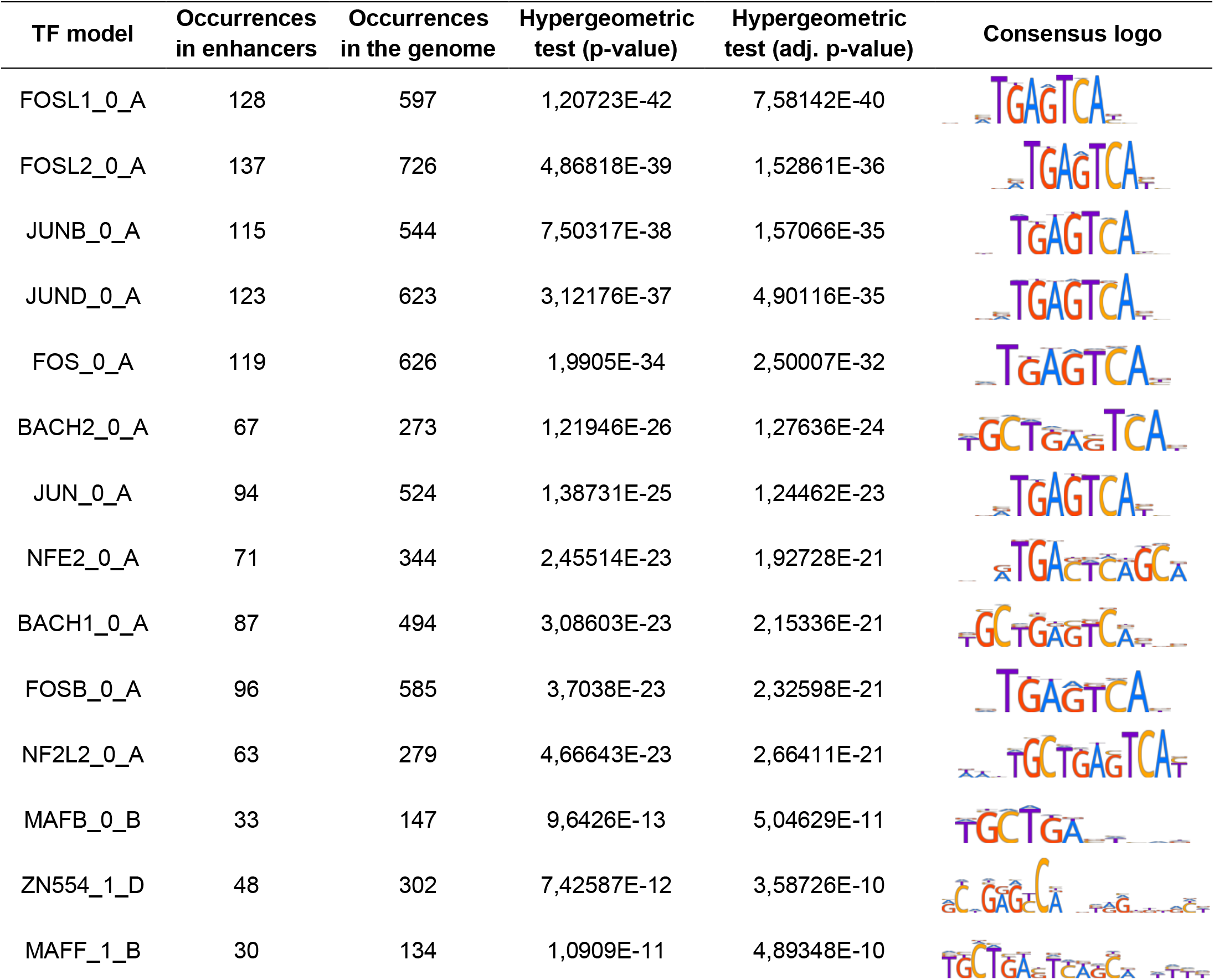
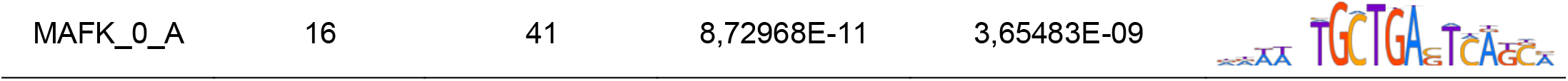
top 15 TF binding probabilities in glioma enhancers calculated by the hypergeometric test. Obtained p-values were corrected using the Benjamini-Hochberg method.

| TF model | Occurrences in enhancers | Occurrences in the genome | Hypergeometric test (p-value) | Hypergeometric test (adj. p-value) | Consensus logo |
| --- | --- | --- | --- | --- | --- |
| FOSL1_0_A | 128 | 597 | 1,20723E-42 | 7,58142E-40 |  |
| FOSL2_0_A | 137 | 726 | 4,86818E-39 | 1,52861E-36 |  |
| JUNB_0_A | 115 | 544 | 7,50317E-38 | 1,57066E-35 |  |
| JUND_0_A | 123 | 623 | 3,12176E-37 | 4,90116E-35 |  |
| FOS_0_A | 119 | 626 | 1,9905E-34 | 2,50007E-32 |  |
| BACH2_0_A | 67 | 273 | 1,21946E-26 | 1,27636E-24 |  |
| JUN_0_A | 94 | 524 | 1,38731E-25 | 1,24462E-23 |  |
| NFE2_0_A | 71 | 344 | 2,45514E-23 | 1,92728E-21 |  |
| BACH1_0_A | 87 | 494 | 3,08603E-23 | 2,15336E-21 |  |
| FOSB_0_A | 96 | 585 | 3,7038E-23 | 2,32598E-21 |  |
| NF2L2_0_A | 63 | 279 | 4,66643E-23 | 2,66411E-21 |  |
| MAFB_0_B | 33 | 147 | 9,6426E-13 | 5,04629E-11 |  |
| ZN554_1_D | 48 | 302 | 7,42587E-12 | 3,58726E-10 |  |
| MAFF_1_B | 30 | 134 | 1,0909E-11 | 4,89348E-10 |  |

The ATAC-seq dataset from LN18 and LN229 glioma cells contained as many as 94 c-Jun motifs (Table 1, Additional file 1: Table S1). The GBM ATAC-seq dataset did not contain all of these motif occurrences (Fig. 4C, GBM1 and GBM2 TFBS predictions). Comparison of the enhancers (H3K27ac signal between GBM and DAs) revealed only few significantly different regions in GBMs (Additional file 1: Table S3). This suggests that the distribution of activating histone marks in the majority of distal regulatory areas is likely to be similar in both DAs and GBMs, or that tumor-derived data is too noisy to detect subtle differences between these tumors. Next, DNA methylation levels of c-Jun motifs and their flanking regions (c-Jun motif +/- 20bp) within enhancers were compared among GII/GIII-IDHwt, GIV and GII/GIII-IDHmut samples from Glioma Atlas [4]. Out of 94 such loci, nine had significantly different DNA methylation levels among glioma groups with the highest median beta values for GII/GIII-IDHmut (FDR<0.05). Within the nine loci we assigned short sequences rich in cytosines: C-rich regions highly overlapping c-Jun motifs. DNA methylation levels of C-rich regions were confirmed to differ significantly among glioma groups (Additional file 2: Fig. S5A) and especially between GII/GIII-IDHwt and GIV with significantly lower levels for GIV, in some cases (Additional file 2: Fig. S5B). This strongly suggests that differential DNA methylation of these nine loci may affect c-Jun motif binding resulting in changed activity of the enhancer.

### c-Jun binds to the *VIM* gene promoter in human glioma cells

Our results suggest that c-Jun is involved in the control of *VIM* expression, which codes for an intermediate filament essential for cell migration, adhesion, and cell division [28,29]. To verify if c-Jun binds to the *VIM* gene promoter, we performed an electrophoretic mobility shift assay (EMSA) using nuclear extracts from normal human astrocytes (NHA), low grade glioma-derived cell cultures (WG12) and two human established glioma cell lines derived from GBM (LN18, LN229). First, we investigated c-Jun expression in these cells. In all tested cells, there was no significant difference in c-Jun mRNA and protein levels (Fig. 5A, B). In EMSA experiments, nuclear extracts from glioma cells bound to the fragment of DNA from the *VIM* promoter, producing a clear shift of the labelled probe (Fig. 5C). The strongest binding to the *VIM* promoter was found in extracts from LN18 and LN229 (Fig. 5C, D) and protein-DNA complexes from three experiments were evaluated by densitometry and quantified (Fig. 5D). Significant differences in the c-Jun binding to the *VIM* promoter were found between GBM and WG12 cells. c-Jun binding to the *VIM* promoter was relatively high in NHA, which may be due to proliferation of these cells in culture conditions (Fig. 5C, D). The findings show that c-Jun binds strongly to the *VIM* promoter in malignant glioma cells. The probe was further shifted after the addition of anti-c-Jun antibody (Fig. 5E), indicating the presence of c-Jun in the DNA-protein complex. The reduction of DNA-protein complexes after the addition of the unlabeled probe competing for c-Jun binding confirmed the binding’s specificity. Next, we used SP600125 (SP) inhibitor of JNK kinases, which phosphorylate c-Jun and studied the impact on *VIM*. We treated LN18 cells with 10 µM concentration of SP inhibitor for 3 h and analyzed the protein levels using Western blot. The results from three experiments were evaluated by densitometry and quantified (Fig. 5F). SP inhibitor efficiently decreased the phosphorylation of S63 c-Jun, which resulted in the significant reduction of the level of VIM. The total c-Jun levels were also affected by the SP treatment, however not significantly, which was possibly due to the inhibition of the positive autoregulatory loop in c-Jun/*JUN* expression [30]. These results show that c-Jun regulates the levels of VIM.

**Fig. 5.**
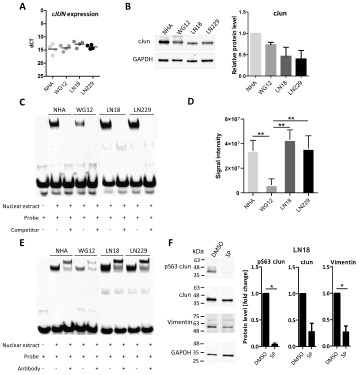
c-Jun transcription factor binds to the *VIM* promoter in human astrocytes and glioblastoma cell lines. c-Jun levels in normal human astrocytes (NHA), low-grade glioma patient derived cell cultures (WG12) and established glioma cell lines (LN18, LN229). **(A)** *c-JUN* mRNA expression was evaluated by RT-qPCR. Data were normalized to the expression of *GAPDH* mRNA determined in the same sample. **(B)** Protein levels of c-Jun analyzed by Western blot and densitometry of immunoblots determined from three experiments, mean ± SD. Data were normalized to the levels of GAPDH in the same sample. The densitometry is presented as relative values to NHA set as 1. P-values were calculated using GraphPad software and considered significant when *p < 0.05 (t-test). **(C)** DNA-binding activity of to double-stranded DNA from the Vimentin promoter site. EMSA was performed using the LightShift Chemiluminescent EMSA Kit. Nuclear extracts were isolated from NHA, WG12, LN18 and LN229. Unlabeled competitor probes were added to lanes 3, 5, 7, and 9. **(D)** Densitometry analysis of EMSA presented as signal intensity of three independent experiments ± SD. One-way ANOVA with Dunnett’s post-hoc test revealed significant differences at **p < 0.01. **(E)** Supershift EMSA assay for measuring c-Jun transcription factor binding to DNA from Vimentin promoter. Antibody against c-Jun was added to lanes 3, 5, 7, and 9 to verify if the observed shift of the probe band in the gel was dependent on c-Jun binding. **(F)** Impact of inhibition of c-Jun phosphorylation on the level of Vimentin. LN18 cells were treated for 3 h with SP600125 (SP) inhibitor of JNK kinases, which phosphorylates c-Jun. Protein levels were analyzed by Western blot and densitometry analysis from three experiments, mean ± SD. The densitometry is presented as relative values to control (cells treated with DMSO, set as 1). Data were normalized to the levels of GAPDH in the same sample. P-values were calculated using GraphPad software and considered significant when *p < 0.05 (t-test).

### DNA methylation at the promoters of c-Jun putative targets differs in low- and high-grade gliomas

DNA methylation may affect binding of a TF to a specific site, therefore we analyzed DNA methylation patterns at the *JUN* promoter and promoters (2 kb upstream/500 bp downstream relative to TSS) of c-Jun putative targets in GII/GIII-IDHwt and GIV glioma patients from the Glioma Atlas [4]. Among the genes potentially controlled by c-Jun, we discovered two clusters: one cluster contains genes with high DNA methylation in GII/GIII-IDHmut and GII/GIII-IDHwt gliomas but low in GIV gliomas (*RIN1, RAB36, UPP1, SLFN12* and *VIM*), and the second cluster encompasses genes with low DNA methylation (beta values ∼0) regardless of a tumor grade (*PTN, FOSL2, FAM111B, SIAH2, SPATA1, TMEM43, TRIB1* and *GPR3*) (Fig. 6A).

**Fig. 6.**
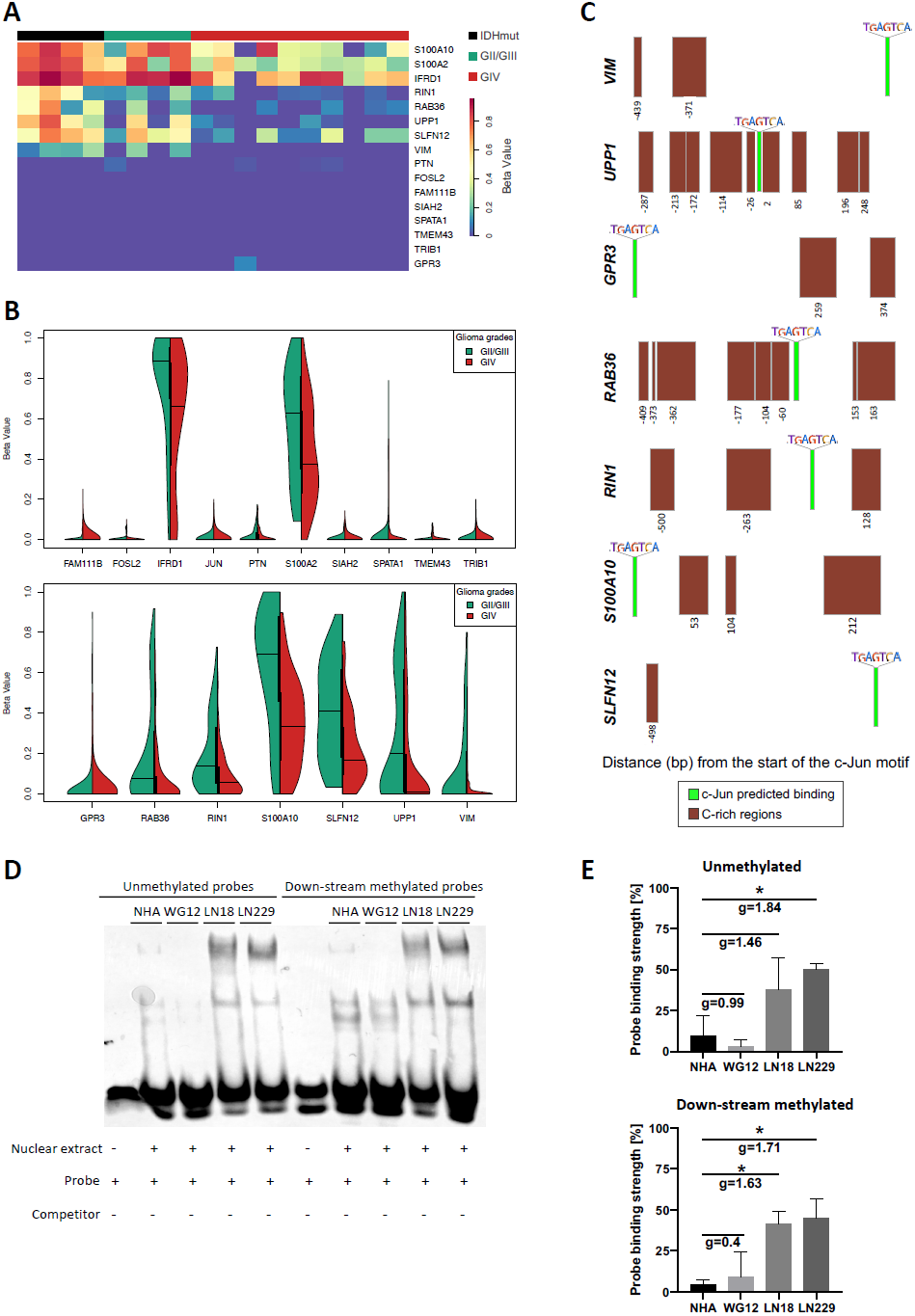
DNA methylation differs in cis-regulatory regions in c-Jun gene targets. **(A)** Heatmap of hierarchical clustering analysis showing median DNA methylation of promoters of c-Jun target genes (2kb upstream and 500bp downstream relative to TSS) in glioma samples. The following labels were used: IDHmut (4 GII/GIII tumors), GII/GIII (4 only GII/GIII-IDHwt tumors) and GIV (10 IDHwt tumors). DNA methylation levels showed as beta values, with 0.0-0.2 representing hypomethylated cytosines and 0.6-1.0 representing hypermethylated cytosines. **(B)** Differences in beta values distribution in gene promoters with the predicted c-Jun TFBS in GII/GIII-IDHwt versus GIV glioma samples that are statistically non-significant (upper panel) and statistically significant (bottom panel). **(C)** Distance of c-Jun motif to the beginning of differentially methylated C-rich regions between high- and low-grade gliomas. Green boxes represent a c-Jun predicted binding site while brown boxes show each C-rich region that was found significantly differently methylated between low- and high-grade glioma samples (Chi-squared test at significance level adjusted p<0.05). **(D)** Competitive electrophoretic mobility assay (EMSA) was used for measuring proteins binding affinity of nuclear extracts from tumor derived cell lines and normal human astrocytes, to the methylated and unmethylated double-stranded DNA from the *UPP1* promoter site. Binding of proteins from NHA, WG12, LN18, and LN229 to the methylated and unmethylated probes was evaluated. Lane 1,6: unlabeled probes; lanes 2-5 and 7-10: protein binding. **(E)** Probe binding strength in EMSA densitometry analysis is expressed as a percentage of the overall variation across all cell lines. Fisher’s LSD tests revealed significant differences at *p<0.05 and **p<0.01; Hedge’s effect size (g) indicates the strength of the difference of the binding between the studied cell lines (medium effects g≈0.5; large effects g≈0.8). The data are presented as the mean SD of three experiments.

Methylation of the c-Jun promoter was low and similar in GII/III-IDHwt and GIV glioma samples (Fig. 6B), suggesting that its differential expression is not regulated by DNA methylation. Many c-Jun target genes had similar DNA methylation at promoters in low- and high-grade gliomas (Fig. 6B, upper panel). The methylation pattern of the promoters of other c-Jun target genes clearly differed (Fig. 6B, bottom panel) with low DNA methylation at their regulatory regions in GIV gliomas (Chi-square test for two independent groups, FDR<0.05), also observed in other types of human cancer [31]. Methylation of a specific cytosine within a c-Jun motif instance (1 cytosine in the c-Jun motif, “dvTGAGTCAYh”, HOCOMOCO version 11) within the promoters of the potential c-Jun targets was found. In several cases, CpG methylation in the flanking regions of predicted c-Jun binding sites varied in gliomas of different WHO grades (Fig. 6C, brown boxes). These results suggest that DNA methylation could control expression of c-Jun regulated genes by affecting TF binding to the regulatory DNA regions.

To validate our previous findings, we explored the TCGA dataset [27]. While the Glioma Atlas and the TCGA datasets had different cytosine coverage, the latter contains more samples and is an independent dataset. We found that the IDH phenotype causes a clear hypermethylated pattern in the majority of the gene promoters (Additional file 2: Fig. S5C). Moreover, many of these genes (*PTN, SIAH2, FAM111B, TMEM43, FOSL2, TRIB1* and *SPATA1*) were hypomethylated regardless of tumor grade (Additional file2: Fig. 5SC). When GII/GIII-IDHmut gliomas were compared to GIV gliomas, significant differences of DNA methylation in the promoters of *S100A2, RAB36, RIN1, UPP1*, and *VIM* were found (Additional file2: Fig. 5SC). Finally, when DNA methylation and transcriptomic data were correlated, DNA methylation was significantly negatively related to gene expression in the majority of cases (Additional file2: Fig. 5SD). The results suggest that methylation of some c-Jun target genes might have an impact on c-Jun binding and transcription of those genes.

### c-Jun binding to the *UPP1* gene promoter is unaffected by DNA methylation

DNA methylation can repress transcription by preventing specific TFs binding to their recognition motifs, while removing site-specific DNA methylation has been shown to reverse the process [32,33]. In gliomas, DNA methylation is tightly associated with mutations in genes coding for the isocitrate dehydrogenase (IDH1/2) which results in glioma CpG island methylator phenotype (G-CIMP) [34,35]. Patients with G-CIMP+ tumors have a better prognosis [36]. Interestingly, specific CpG loci in genes of the homeobox family (*HOXD8, HOXD13 and HOXC4*), among other regulators are differentially hypermethylated between patients with short- and long-term survival [36].

We explored the previously collected data on DNA methylation in gliomas of different grades [4]. The analysis of DNA methylation profiles in the promoters of GIV-specific genes with c-Jun binding sites showed different levels of methylated cytosines within those promoters (Fig. 6A). Furthermore, some proximal c-Jun binding sites were differently methylated in C-rich regions (Fig. 6C). In GII/GIII-IDHwt gliomas proximal areas to c-Jun binding (−26 bp and +2 bp) exhibited highly methylated cytosines in comparison to similar sites in GBMs (Fig. 6C). In particular, the *UPP1* gene promoter contained numerous differentially methylated cytosines around the c-Jun binding site.

To determine the effects of DNA methylation on c-Jun binding, we determined binding of the nuclear extracts from NHA, WG12, LN18 and LN229 glioma cells to unmethylated and methylated *UPP1* probes using EMSA. Nuclear extracts from LN18 and LN229 produced a shift of the *UPP1* probe and the binding of nuclear extracts from WG12 cells to both probes was lower in comparison to LN18 and LN229 cells (Fig. 6D). Although the binding to the unmethylated/methylated probes was noticeably stronger in malignant glioma cells (LN18 and LN229) when compared with normal astrocytes and WG12 cells (Fig. 6D-E), the binding to the *UPP1* unmethylated probe and to the methylated probe in malignant glioma cells remained similar (Fig. 6E). To clarify this issue, EMSA experiments were carried out separately with nuclear extracts from each of the cell lines. Only in NHA, methylation of the probes in the flanking regions of the *UPP1* gene promoter had a small effect on the transcription factor binding (Additional file 2: Fig. S6).

## Discussion

In this study, we analyzed a broad range of publicly available and experimental data generated to identify novel TF-gene regulatory networks contributing to transcriptional deregulation in malignant brain tumors. Using chromatin accessibility data from tumor samples and cultured cells, we identified GBM-specific TFs binding sites that were also present in human LN18 and LN229 glioma cells. The majority of accessible chromatin regions identified by ATAC-seq were localized in the promoters. Some TFBS, such as binding sites for AP2D, PAX5, and ZFX, were highly enriched within open chromatin areas (Fig. 1C). The transcription factor activator protein 2 (AP-2) is involved in a variety of pathological and carcinogenic processes [37]. Transcription factor PAX5 (*Paired Box Gene 5*) promotes gliomagenesis and cooperates with factors such as MYC, FOS, or JUN which are highly expressed in gliomas [38]. The zinc finger and X-linked transcription factor (ZFX) is associated with proliferation, tumorigenesis, and patient survival in a variety of human cancers [39] and maintains self-renewal and tumorigenic potential of glioma stem like cells by upregulating *c-Myc* expression [40]. While the available data do not allow to solve a “chicken-egg” dilemma, it is tempting to speculate that increased chromatin accessibility in GBMs and the enrichment of specific TFBS in the promoters of cancer-related genes, result in the establishment of novel TF-gene regulatory network driving tumorigenesis.

A close inspection of genes overexpressed in GBMs versus benign gliomas revealed the specific TFBS enrichment in the promoters of these genes which is indicative of the contribution of a specific TF to glioma development or progression. c-Jun appeared as a prominent candidate for novel TF-gene regulatory network driving tumorigenesis (Fig. 1D). It is a well-known proto-oncogene, involved in proliferation, angiogenesis, migration and apoptosis in several cancers [41,42]. Interestingly, we found that in the majority of human cancers (bladder, breast, skin, cervical/endocervical, ovarian, lung and uterus), *JUN* expression is higher in normal tissues than in tumors, but only in thymomas and GBMs, *JUN* was significantly overexpressed in tumors when compared to neighboring tissues (Fig. 2A). Moreover, high-grade gliomas showed increased *JUN* expression (Fig. 2B).

We detected c-Jun binding motifs in the promoters of 16 genes overexpressed in GBMs. Many of these genes (Fig. 2C-E), such as *VIM, FOSL2, PTN, GPR3, SIAH2, UPP1, S100A2* are associated with cell migration and endothelial-mesenchymal transition (EMT) in several cancers [43–50]. The positive correlation between c-Jun mRNA/protein and target gene expression indicates that c-Jun most likely regulates the predicted targets in GBMs (Fig. 3D-E). All these genes accurately predicted patient survival in LGGs but not GBMs (Additional file 2: Fig. S3-S4). One explanation for the observed lack of association in GBMs could be that the expression of these genes is very high in most of GBMs, making the distinction between low-expressing and high-expressing patients difficult.

The fact that several putative Jun-(JunB, JunD) and fos-related factors (c-Fos, FRA1, FRA2, and FosB) TFBS overlap with c-Jun binding sites (Fig. 2E) envisions the complexity of c-Jun-regulated networks. c-Jun (which, along with those TFs, forms the AP-1 complex) may interact with different TF forming homo- or heterodimers [51] at the specific promoters and differentially regulate target gene expression. In the AP-1 complex, a selection of a dimerizing partner influences not only the DNA recognition properties but also the regulatory function of a given TF as some JUN family members lacks transactivating domains [52]. In glioma enhancers with the H3K27ac activating histone marks, we identified the enrichment in c-Jun binding motifs together with motifs for other TFs (Table 1, Fig. 4C) supporting a role of c-Jun in driving GBM specific expression. Expression of genes coding for GBM-specific TFs was higher in malignant gliomas (Fig. 3B, upper panel), as is exemplified by the expression of *HOX* genes, *JUN* and *TWIST* according to previous reports [53,54].

Overexpression *of VIM* as measured by immunohistochemistry is a poor prognostic factor in GBMs [28]. We predicted that c-Jun may drive high *VIM* expression in GBMs and verified this prediction in human GBM cells where c-Jun binds to the *VIM* gene promoter, as demonstrated by EMSA and supershift assays (Fig. 5C-E). GBM-derived cells had more DNA-protein complexes than cells derived from the benign GII glioma (Fig. 5D). Because the concentrations of nuclear proteins used for reactions were the same across all cell lines with equivalent levels of c-Jun (Fig. 5A-B), the lower binding suggests that transcription of c-Jun target genes is less effectively activated in those cells. Regulation of VIM expression by c-Jun in glioma cells was confirmed with the use of a JNK inhibitor, which decreased phosphorylated c-Jun and VIM levels (Fig. 5F). This is consistent with previous findings in epithelial cells [55], indicating that c-Jun plays a widespread role in oncogenic processes in various cancer types. The physical access of TFs to DNA can be modulated by nucleosome positioning, histone modifications or DNA methylation [33,56]. When we examined DNA methylation levels at the promoters of c-Jun and its targets in low- and high-grade gliomas, we found differences in DNA methylation in putative c-Jun target genes *S100A10, S100A2, IFRD1, RIN1, RAB36, UPP1, SLFN12* and *VIM* (Fig. 6A-B). DNA methylation differences patterns varied primarily in the flanking regions than at c-Jun binding sites (Fig. 6C). Although we found distinct c-Jun binding in the *UPP1* gene promoter when comparing various glioma cell lines by examining DNA-protein complexes (Fig. 6C), its binding to the *UPP1* unmethylated probe and to the methylated probe in malignant glioma cells remained similar (Additional file 2: Fig. S6), indicating that flanking hypermethylated regions may not have an impact on c-Jun binding to the specific binding site.

## Conclusion

Understanding particular TF-gene regulatory networks responsible for the dysregulated expression of certain genes in malignant gliomas holds a key to potential pharmacological manipulations. We demonstrate the increased expression of *JUN* mRNA/protein in GBMs (which is uncommon in other pan-cancer genomes) and the enrichment of c-Jun binding motifs in the accessible chromatin of GBM-specific genes. These genes are implicated in tumorigenesis, and high expression of putative targets of c-Jun is associated with poor survival of glioma patients. We demonstrate with biochemical methods the stronger binding of c-Jun to *VIM* and *UPP1* promoters in human malignant glioma cells. The inhibition of phosphorylation of c-Jun resulted in reduced levels of VIM, indicating that c-Jun is involved in the regulation of VIM levels.

## Methods

### Sample collection and human glioma cell cultures

Human established glioblastoma LN18 and LN229 cells were obtained from the American Type Culture Collection (ATCC). The cells were cultured in Dulbecco’s Modified Eagle Medium (DMEM) supplemented with 10% fetal bovine serum (FBS, ThermoFisher Scientific), 100 units/mL penicillin and 100 μg/mL streptomycin. In some experiments LN18 were treated with 10 µM SP600125 inhibitor (Biotechne, cat. no 1496/10) for 3 h, as a control cells were treated with 0.05% DMSO. Freshly resected glioma specimens were acquired from two neurosurgical departments of the Medical University of Warsaw and the Mazovian Brodno Hospital. The tissue collection protocol was approved as described [57]. Tumor samples were transported in DMEM/F-12 medium on ice and processed immediately after surgical resection. Tumor samples were transferred to cold PBS, minced with sterile scissors or a scalpel on a Petri dish kept on ice, and then homogenized with a chilled manual glass mince [57]. Each patient signed a written consent form for the use of tumor tissues, and the samples were then anonymized. Human glioma LN18, LN229 cell lines were from American Type Culture Collection (ATCC, Manassas, VA, USA); GBM patient-derived glioma primary cultures WG12 were generated and cultured as described [58] in DMEM/Nutrient Mixture F-12, GlutaMAX™ medium, supplemented with 10% FBS (Gibco Life Technologies, Rockville, MD, USA) and antibiotics. See Additional file 2: Fig. S7 for WG12 cell line’s detailed characterization. Normal human astrocytes (NHA cat # CC-2565, Lonza Walkersville, MD, USA) were cultured in a commercial Astrocyte Growth Medium (AGM™ cat # CC-3186, Lonza Walkersville, MD, USA) supplemented with 3% FBS, 1% L-glutamine, 0.1% ascorbic acid, 0.1% human EGF, 0.1% gentamicin, and 0.0025% recombinant human insulin. NTERA-2 cl.D1 (cat # CRL-1973 ™) were purchased from ATCC (Manassas, VA, USA) and cultured in DMEM with GlutaMax-1 and supplemented with 10% FBS. The L0125, and L0627 GBM GSCs (glioma stem-like cells) were provided by Dr Rossella Galli (San Raffaele Scientific Institute, Milan, Italy) [59–61]. Spheres L0125 and L0627 were expanded *in vitro* in serum-free medium supplemented with 2% B27, 20 ng/ml rh EGF and 20 ng/ml rh bFGF as described before [62,63]. For differentiation experiments, spheres were triturated for a single cell suspension and seeded onto laminin-coated plates in the medium without cytokines (rh EGF and rh bFGF) but containing 2% FBS and were incubated for 7 days [62]. All cell cultures were grown in a humidified atmosphere of CO2/air (5%/95%) at 37 °C.

### DNA isolation

Genomic DNA was extracted from 50-100 mg of glioma sample (depending on a size of the original tumor specimen) and from 2×10^7^ LN18 and LN229 cells using Tri-Reagent (Sigma-Aldrich, Munich, Germany). DNA purity was estimated using NanoDrop 2000 (Thermo Scientific, NanoDrop products, Wilmington, USA) and quantity with Agilent Bioanalyzer as described [4].

### ATAC-sequencing

Tumor sample aliquots corresponding to 50-100 mg of tissue were drawn through a syringe needle approximately 50 times. Mechanical homogenization was followed by 5 minutes of centrifugation at 2,400g at 4°C. Each pellet was resuspended in 10 ml of cold lysis buffer L1 (50 mM HEPES KOH, pH 7.5, 140 mM NaCl, 1 mM EDTA pH 8.0, 10% glycerol, 5% NP-40, 0.25 %Triton X-100, proteinase inhibitor cocktail) and shaken for 20 minutes at 4°C. The tissue was then mechanically disrupted, residual debris pre-cleared by filtration through nylon mesh filters, and the lysis buffer was replaced with PBS. Cells were automatically counted using the NucleoCounter NC-100, and 50,000 cells were lysed as previously described [4]. The extracts were filtered using Zymo DNA Clean and Concentrator 5 columns. ATAC-seq library preparation was carried out as described [4]. Finally, ATAC-seq libraries were visualized on a Bioanalyzer 2100 (Agilent Technologies, Santa Clara, CA) and DNA concentration was estimated. Libraries were run in the Rapid Run flow cell and paired-end sequenced (2×76 bp) with the HiSeq 1500 (Illumina, San Diego, CA 92122 USA).

### Processing of ATAC-sequencing data

The FastQC tool was used to evaluate a quality of raw fastq data [64]. After trimming ATAC-seq reads with the FASTQ trimmer [65], only reads with a quality of 10 or higher were considered. Reads with incorrect pairing and a length of less than 20 bp were discarded. Using the default parameters, the Bowtie2 aligner [66] was used to map the reads to the human genome (hg38). Only high-quality reads (MAPQ > 30), correctly paired read mates, and uniquely mapped reads were considered for downstream analysis. PicardTools [67] was also used to find and eliminate PCR duplicates. The following parameters in MACS2 were used to center a 200 bp window on the Tn5 binding site (5’ ends of reads represent the cut sites), which is more accurate for ATAC-seq peaks: *--broad --nomodel --shift −100 --extsize 200*. Resulting peaks were then intersected with human (hg38) ENCODE blacklisted genomic regions (https://github.com/Boyle-Lab/Blacklist/tree/master/lists) to eliminate anomalous and unstructured signals from the sequencing.

### Selection of genes differentially expressed in gliomas of different WHO grades

We used TCGA data to find overexpressed genes in glioblastomas (GIV) compared to benign gliomas (GII, WHO grade II) using normalized RNA-seq expression values (Fragments Per Kilobase of transcript per Million mapped reads, FPKM). RNA-seq data from 408 glioma patients (248 GII gliomas and 160 GIV gliomas) were analyzed. The biotype of genes from RNA-seq data was restricted to protein-coding genes [68]. We randomly sampled 20 GII and 20 GIV patients from the normalized count matrix (n=200 times) to maximize statistical power and robustness of the gene selection. The sample function from the base R library (version 3.6.2) was used as a sampling technique, with each sample having an equal chance of being chosen.

We then calculated Student’s t-test p-values for all the genes for each of 200 random comparison of 20 GII vs 20 GIV, the p-values were then corrected using multiple testing (FDR), and the means of these adjusted-p values from all of these comparisons were calculated. We used DESeq2 methods [69] on the same TCGA dataset to find changes in gene expression and determined the directionality of the change using the log fold change (logFC) criteria. Overexpressed genes in GBM were those that changed significantly based on FDR and had a positive logFC. To investigate the biological significance of our gene selection, we performed pathway enrichment analysis with the ClusterProfiler [70] R library using the Gene Ontology (Biological Processes) and KEGG databases. Here, only genes that differed significantly between glioma grades (GIV vs. GII) were retained (adjusted p-value means < 0.01).

### Prediction of transcription factor binding from ATAC-seq data

The human HOCOMOCO v11 motif database [20] in the MEME motif format was used to find TFs that could potentially bind to promoters within open chromatin regions. Using the FIMO tool [71], position weight matrices (PWMs) were used to scan the FASTA file of the human genome (hg38). The background nucleotide frequency from hg38 was used and all motif occurrences with a p-value less than 1e^-4^ on both DNA strands were considered. Motifs found on the mitochondrial genome were discarded for the subsequent analysis. Overall, motif occurrences were computed independently for each of the 735 motifs. The BMO algorithm [72] was used to classify TF binding in human glioma cells and human glioblastoma samples. For further analysis, only motif instances expected to be bound with adjusted p-values below 0.05 (Benjamini-Yekutieli correction procedure) were used. Only motif instances at the same chromosomal localization in LN18 and LN229 human glioblastoma cells were considered. We intersected the resulting TFBS with the promoters of protein coding genes after selecting motif instances that were common in both glioma cell lines. Transcription start sites and their flanking DNA regions upstream (1.5kb) and downstream (1.5kb) were used to identify gene promoters [68]. If a particular TF was predicted to bind twice in a single promoter, it was counted as one TFBS because we only considered one TFBS per promoter. By focusing on the top TFs, we could determine the importance of a specific TF and its relationship to gene dysregulation. Finally, the BMO results from two samples of human glioblastoma were compared to the TFBS found in the LN18 and LN229 glioblastoma cells.

### RNA isolation from human glioma cells and RNA-seq processing

Total RNA from glioma cells was isolated suing a Qiagen RNeasy kit. In brief, 1×10^6^ cells were lysed with 350 µL RLT buffer supplemented with 1% β-mercaptoethanol. The extraction procedure was carried out in accordance with the manufacturer’s instructions. The total RNA was eluted with 25 µl of sterile H_2_O and its concentration was estimated with Nanodrop 2000 (Thermo Scientific, NanoDrop products, Wilmington, USA). The KAPA Stranded mRNA Sample Preparation Kit was used to prepare polyA enriched RNA libraries (Kapa Biosystems, Wilmington, MA, USA). Trimmomatic [73] (version 0.36) with default parameters was used for the transcriptomic analysis to remove Illumina adapters and low-quality reads. Then, RNA sequencing reads were aligned to a reference genome sequence (hg38) with the twopassMode Basic choice enabled in STAR aligner [74] (version 2.6) and all other parameters were set to default. Only properly oriented pairs of reads were considered for downstream analysis. Flag read duplicates and optical duplication estimation was done using MarkDuplicates from Picard Tools [67] (version 2.17.1). RNA-seq mapped reads in paired and reverse stranded mode were summarized and counted by genes using featureCounts software [75] (version 1.5.3). Only genes that were uniquely mapped and had MAPQ mapping quality values of 255 were considered. Raw counts from featureCounts were converted to FPKM values, and genes encoding various transcription factors were selected for further investigation.

### Gene expression profiling in pan-cancer and paired normal tissues

We used the Gene Expression Profiling Interactive Analysis (GEPIA2) [76] to determine *JUN*’s expression in various human cancers, including in brain tumors, Transcripts per million (TPM) values were extracted from different TCGA and Genotype-Tissue Expression (GTEx) datasets, and median gene expression in each cancer and paired normal tissues was calculated and used as an input to R. The following cancers and corresponding healthy tissues were examined: Adrenocortical carcinoma (ACC) and Adrenal Gland; Bladder Urothelial Carcinoma (BCLA) and bladder; Breast invasive carcinoma (BRCA) and breast; Cervical squamous cell carcinoma and endocervical adenocarcinoma (CESC) and cervix uteri; Colon adenocarcinoma (COAD) and colon; Lymphoid Neoplasm Diffuse Large B-cell Lymphoma (DLBC) and blood, Esophageal carcinoma (ESCA) and esophagus; Glioblastoma multiforme (GBM) and brain; Kidney Chromophobe (KICH) and kidney; Kidney renal clear cell carcinoma (KIRC) and kidney; Kidney renal papillary cell carcinoma (KIRP) and kidney; Acute Myeloid Leukemia (LAML) and bone marrow; Brain Lower Grade Glioma (LGG) and brain; Liver hepatocellular carcinoma (LIHC) and liver; Lung adenocarcinoma (LUAD) and lung; Lung squamous cell carcinoma (LUSC) and lung; Ovarian serous cystadenocarcinoma (OV) and ovary; Pancreatic adenocarcinoma (PAAD) and pancreas; Prostate adenocarcinoma (PRAD) Prostate, Rectum adenocarcinoma (READ) and colon; Skin Cutaneous Melanoma (SKCM) and skin; Stomach adenocarcinoma (STAD) and stomach; Testicular Germ Cell Tumors (TGCT) and testis; Thyroid carcinoma (THCA) and thyroid; Thymoma (THYM) and blood; Uterine Corpus Endometrial Carcinoma (UCEC) and uterus; Uterine Carcinosarcoma (UCS) and uterus. The *JUN* mRNA expression profile was compared between tumor samples (TCGA) and paired normal tissues (TCGA normal + GTEx normal), and statistical significance was determined using one-way ANOVA and disease state (Tumor versus healthy tissue of tumor origin).

### ChIP-sequencing

The QIAseq Ultra Low Input Library Kit was used to create DNA libraries for chromatin immunoprecipitation with the appropriate antibodies (QIAGEN, Hilden, Germany), as described [4]. End-repair DNA was used, adenosines were added to the 3′ ends of dsDNA to create “sticky-ends”, and adapters (NEB, Ipswich, MA, USA) were ligated. Following adapter ligation, uracil was digested in an adapter loop structure by USER enzyme from NEB (Ipswich, MA, USA). Using NEB starters, adapters containing DNA fragments were amplified by PCR (Ipswich MA, USA). The Agilent 2100 Bioanalyzer with the Agilent DNA High Sensitivity chip was used to evaluate the library’s quality (Agilent Technologies, Ltd.) To quantify and evaluate the obtained samples, the Nanodrop spectrophotometer (Thermo Scientific, NanoDrop products, Wilmington, USA), Quantus fluorometer (Promega Corporation, Madison, USA), and 2100 Bioanalyzer were used to quantify and evaluate the obtained samples (Agilent Technologies, Santa Clara, USA). The average library size was 300 bp. Libraries were run in the rapid run flow cell and were single-end sequenced (65 bp) in the rapid run flow cell on HiSeq 1500 (Illumina, San Diego, CA 92122 USA).

### Comparison of H3K27ac histone modification across glioma grades

We had acquired histone ChIP-seq data from gliomas of different grades from the Glioma Atlas [4] focusing on activated enhancers from eight diffuse astrocytomas (DAs) and ten GBMs. We used the DESeq2 method [69] to identify H3K27ac ChIP-seq signal differences within enhancer peaks to better capture the differences in active enhancer marks between glioma grades. First, we filtered out peaks found in only one tumor sample and the resulting peakset was used to count single-end reads from BAM files using the featureCounts [75] tool. H3K27ac signal differences between GBMs and DAs were identified and only regions with adjusted p-values 0.05 were considered as statistically significant.

### Annotation of glioma enhancers and their association with TFBS

To select active enhancers in glioma, we considered the presence of H3K27ac peaks in non-promoter regions and used a set of active enhancers identified previously [4]. First, we used the ChIPseeker (version 1.28.3) library’s peakAnnotation function [77] to pre-filter potential H3K27ac peaks near TSS regions. The resulting set of glioma enhancers was then intersected by chromosomal coordinates with predicted TFBS in glioma cell lines using the tidygenomics R library [78] genome intersection function (version 0.1.2). Furthermore, we performed an integrative analysis of TFBS motifs in enhancers to model the relationship between each TF and distal-regulatory regions. Using the phyper function in R, we calculated probabilities based on the cumulative distribution function of the hypergeometric distribution; the probability of finding the observed number of motif instances within glioma enhancers is represented by the p-values obtained for each of the TF regulatory networks.

### DNA methylation sequencing

EZ DNA Methylation-Lightning Kit was used to bisulfite-convert DNA samples (Zymo Research, Irvine, CA, USA). SeqCap Epi CpGiant Enrichment Kit (Hoffmann-La Roche, Basel, Switzerland) probes were used to enrich each Bisulfite-Converted Sample Library in the predetermined distinct genomic regions of 80.5 Mb capture size, which included 5.6 million CpG sites on both DNA strand. The libraries were created using the “NimbleGen SeqCap Epi Library Workshop Protocol, v1.0” and “SeqCap Epi Enrichment System User’s Guide, v1.2” from Hoffmann-La Roche. In brief, genomic DNA concentration was determined using a Quantus Fluorometer with QuantiFluor dsDNA System (Promega, Madison, WI, USA), and 1 g/mL streptomycin input DNA, as well as 165 pg Bisulfite-Conversion Control (viral unmethylated gDNA; SeqCap Epi Accessory Kit; Hoffmann-La Roche) were fragmented to an average size of 200 20 bp using the Covaris M220 (Covaris, Inc., Woburn, MA, USA). DNA fragments were tested on a 2100 Bioanalyzer using the High Sensitivity DNA Kit (Agilent Technologies, Inc., Santa Clara, CA, USA). Using the KAPA LTP Library Preparation Kit (KAPA Biosystems, Wilmington, USA), SeqCap Adapter Kit A and B (Hoffmann-La Roche), and DNA purification beads, the DNA fragments were “End-Repaired,” “A-tailed,” and the index adapters were ligated (Agencourt AMPure XP Beads; SeqCap EZ Pure Capture Bead Kit; Hoffmann-La Roche). Following that, adapter-enhanced DNA fragments were size-selected using Agencourt AMPure XP beads (SeqCap EZ Pure Capture Bead Kit) and Solid Phase Reversible Immobilization technology to exclude DNA fragments larger than 450 and smaller than 250 bp. Libraries were bisulfite transformed as described [4]. The size of the collected DNA fragments was determined using a 2100 Bioanalyzer and the High Sensitivity DNA Kit (Agilent Technologies, Inc.). Libraries were run in the Rapid Run flow cell and sequenced with paired-end sequencing (2×76 bp) on HiSeq 1500 (Illumina, San Diego, CA 92122 USA).

### Analysis of DNA methylation in published glioma datasets

The methylation analysis workflow was carried out using the CytoMeth tool [79], which takes Fastq files as inputs and returns the calculated DNA methylation levels (beta-values) at a base-pair level. This automated workflow includes: FastQC [64] to assess read quality, BSMAP [80] to map reads to the hg38 reference genome, Picard Tools [67] to remove PCR duplicates and methratio.py to assess coverage statistics and assign methylation levels returned as beta-values. The minimal bisulfite conversion was set to ∼99%. The cytosines in CpG and non-CpG contexts with at least ten reads of coverage were further examined. The analysis was performed on various glioma samples: GII/GIII-IDHwt (n=4), GIV (n=10) and GII/GIII-IDHmut gliomas (n=4). In the end, each sample yielded ∼3.5 × 10^6^ of well-covered cytosines. However, due to DNA degeneration, the total number of cytosines shared by all samples was only ∼350,000. The further analysis focused on differentially methylated regions rather than individual cytosines.

The DiffMeth [81] module was used to compare DNA methylation levels within promoter regions (2 kb upstream/500 bp downstream relative to TSS) as well as c-Jun motif containing regulatory regions within enhancers (50 bp long regions: 10 bp c-Jun motifs extended by flanking regions +/- 20 bp). A standard Chi2 statistical test was used to analyze statistical significance, and all groups were compared to one another. The Chi2 test compared the distribution of beta values reflecting hypo-, medium-, and hyper-methylated cytosines: [0.0-0.2], (0.2-0.6], and (0.6-1.0]; the obtained p-values were corrected with FDR (insignificant if <0.05). DiffMeth [81] was set to detect short regions of similar length to TFBS, with a median length of 22 bp. The only results that were reported were those for methylated cytosines in the CpG context.

### Analysis of DNA methylation in the TCGA dataset

CpG sites in regulatory regions of c-Jun putative target genes were intersected with promoter regions defined as TSS±1 kb with the annotation for the hg19 human genome [82]. GBM and LGG TCGA 450k DNA methylation datasets were downloaded from [27]. For each defined promoter region, the median beta-values of DNA methylation were calculated per each sample. FPKM-normalized TCGA data were uploaded, and Pearson correlation was calculated for selected genes for samples with matching DNA methylation and RNA-seq data. Furthermore, we searched the TCGA for information on DNA methylation in the enhancers and used the available CpG (cg02258482, cg12155676 and cg08003402).

### Survival analyses

The GlioVis web application [83] was used to conduct the analysis on the TCGA data (GBM and LGG datasets). Based on the expression of c-Jun target genes, patients were split into two subgroups (high mRNA and low mRNA levels). The association between c-Jun target expression levels and patient survival was tested using log-rank tests. For each of those genes, Kaplan-Maier plots were generated, and data from censored patients was used in the analyses.

### Immunoblotting and RT-qPCR

Whole-cell protein extracts were prepared, resolved by electrophoresis and transferred to a nitrocellulose membrane (GE Healthcare cat. number 10600003) as described [84]. After blocking with 5% non-fat milk in TBST (Tris-buffered solution pH 7.6, 0.01% Tween-20) the membranes were incubated overnight with primary antibodies diluted in TBST with 5% bovine serum albumin (BSA). The primary antibody reaction was followed by 1 h incubation with horseradish peroxidase-conjugated secondary antibodies. Immunocomplexes were detected using an enhanced chemiluminescence detection system (ECL) and Chemidoc (Biorad). The molecular weight of proteins was estimated with Cozy prestained protein ladder (High Qu GmbH cat. number PRL0102c1). Band intensities were measured by a densitometric analysis of immunoblots with BioRad Image Lab software. P values were calculated using GraphPad software and considered significant when *P < 0.05 (column statistics t-test). Gene expression analysis was performed as described in [4]. Antibodies for Western Blot and the sequences of the used primers are listed in Additional file 1: Table S4.

### Isolation of nuclear extracts and Electrophoretic Mobility Shift Analysis

Nuclear extracts were prepared from cultured cells using a nuclear extraction kit: NE-PER Nuclear and Cytoplasmic Extraction Reagents (Thermo Scientific cat# 78833) according to the manufacturer instructions. Protein concentration was measured using THERMO Labsystems Multiscan EX at wavelength 570 nm with a Bradford Reagent (Sigma Life Science cat no. B6916) and a bovine serum albumin standard (Thermo Scientific cat no. 23209) for calibration. For EMSA, biotin-labelled and unlabeled oligonucleotides were provided by Metabion (Additional file 1: Table S4). For annealing, oligos were dissolved in water, heated to 90°C and let to anneal for 30 min. EMSA was performed using the LightShift Chemiluminescent EMSA Kit (Thermo Scientific cat. #20148) according to the manufacturer instructions. The reactions contained: 40 fmol dsDNA, 5 µg of protein nuclear extracts and 10 mM Tris pH 7.5 buffer with 50 mM KCl, 1 mM DTT, 1.5 mM MgCl _2_, and 1.5 µg Poly (dI-dC) (in 30 μL) and all components were incubated for 30 min at room temperature and subjected to electrophoresis (70 V, 8°C) in 6% polyacrylamide gels with 10% glycerol and Tris– borate–EDTA buffer. Then, electrophoretically separated material was transferred onto a 0.45 µm Biodyne nylon membrane (Thermo Scientific cat. # 77016) in Tris–borate–EDTA buffer and detected by chemiluminescence using a Chemidoc camera (Bio-Rad). For a competition assay, the mixture was pre-incubated with 100-fold unlabeled probe. For a supershift assay, the protein extracts in the reaction mixture were pre-incubated for 30 min with 2 µg of anti-pS63 c-Jun antibody before adding to the DNA. Antibodies and probes used for EMSA are listed in Additional file 1: Table S4.

### Immunocytochemistry

The WG12, L0125, L0627, NTERA cells were seeded onto glass coverslip. L0125 and L0627 cells were differentiated by growing in the presence of 2% FBS. At the appropriate time cells were fixed with 4% PFA pH 7.2, washed, permeabilized with either 0.1% (cytoplasmic staining) or 0.5% (nuclear staining) Triton-X100 and blocked-in mix of 2% donkey serum and 1.5% FBS, followed by 2 h incubation with primary antibodies. Cells were then washed in PBS, incubated with corresponding Alexa Fluor A555 secondary antibodies, counterstained with DAPI (Sigma, 0.001 mg/mL, PBS) and mounted. For reagent specifications, catalogue numbers and concentrations, see Additional file 1: Table S4. Images were taken using the Leica DM4000B fluorescence microscope.

### Bisulfite DNA conversion and methylation-specific polymerase chain reaction (MS-PCR)

DNA was extracted using standard phenol/chloroform methods. The purity and concentration of DNA were estimated after collecting absorbance readings at 260/280 nm. DNA (2 μg) was treated with bisulfite (EpiTect Bisulfite Kit, Qiagen, Hilden, Germany). The modified DNA was amplified using primers specific for methylated or unmethylated MGMT gene promoters, as listed in Additional file 1: Table S4. Each PCR mixture contained 1 μl of DNA, 500 nM of primers, 1x reaction buffer containing 1.5mM MgCl2, and 1 U HotStarTaq DNA Polymerase and 250mM dNTPs (Promega, USA). PCR was performed with thermal conditions as follows: 95 °C for 10 min, 45 cycles of 95 °C for 30 s, 57 °C for 30 s and 72 °C for 30 s with a final extension of 72 °C for 10 minutes. PCR products were visualized using Agilent Tape Station system (Agilent Technologies, Palo Alto, CA, USA), yielding a band of 81 bp for a methylated product and 93 bp for an unmethylated product. Positive methylated and positive unmethylated controls (EpiTect PCR Control DNA Set Qiagen, Hilden, Germany) were included.

## Declarations

### Ethics declarations

This study was approved by The Bioethics Committees of Andrzej Frycz Modrzewski Cracow University, St. Raphael Hospital, Cracow, Poland (Nr. 73/KBL/OIL/2015); Medical University of Silesia, Sosnowiec, Poland; Mazovian Brodno Hospital, Warsaw, Poland (Nr. KNW/0022/KB1/46/I/16).

### Consent for publication

Patients signed an informed consent for the use of their biological material for research purposes.

### Availability of data and materials

The datasets generated and/or analyzed during the current study are available in the Synapse repository [85]. Request to access raw data is available through the European Nucleotide Archive with the accession code ERP125425 and the Gene Omnibus GSE206357. All of the scripts used to generate the computational results presented in this paper can be found in [86].

### Author contributions

Conception and design: A-J.R., B.W. Computational analyses: A-J.R., B.W., M.J.D., cell cultures and biochemical analyses: P.S., K.P., A.E.M., K.W., I.C, K.S. Data interpretation: A-J.R., B.W., M.J.D., B.K., Manuscript writing: A-J.R., B.W., K.P., A.E.M., B.K. Final approval of manuscript: All authors. Accountable for all aspects of the study: All authors.

## Acknowledgments

We thank both the physicians who performed the surgeries and the patients for their consent for use of their biological material for this research.

## Supplementary Information

Additional file 1: Comprises supplemental tables S1-S5 listing search results, the characterization of a grade II glioma cell line (WG12) and the antibodies and primers used in western blot and immunostaining.

Additional file 2: Comprises supplemental figures S1-S7 illustrating the study’s experimental and computational design, transcriptomic analyses from the TCGA between low- and grade-glioma, patient survival analyses concerning c-Jun gene targets, the DNA methylation external validation (TCGA) in c-Jun regulated genes in human gliomas, DNA methylation levels validation in Glioma Atlas concerning c-Jun motifs in putative glioma enhancers, EMSA for nuclear extracts employing probes with different methylation patterns within the *UPP1* promoter, as well as the characterization of a patient-derived primary WG12 cell line.

