## Additional_file_2 for "Regulatory networks driving expression of genes critical for glioblastoma are controlled by the transcription factor c-Jun and the pre-existing epigenetic modifications"

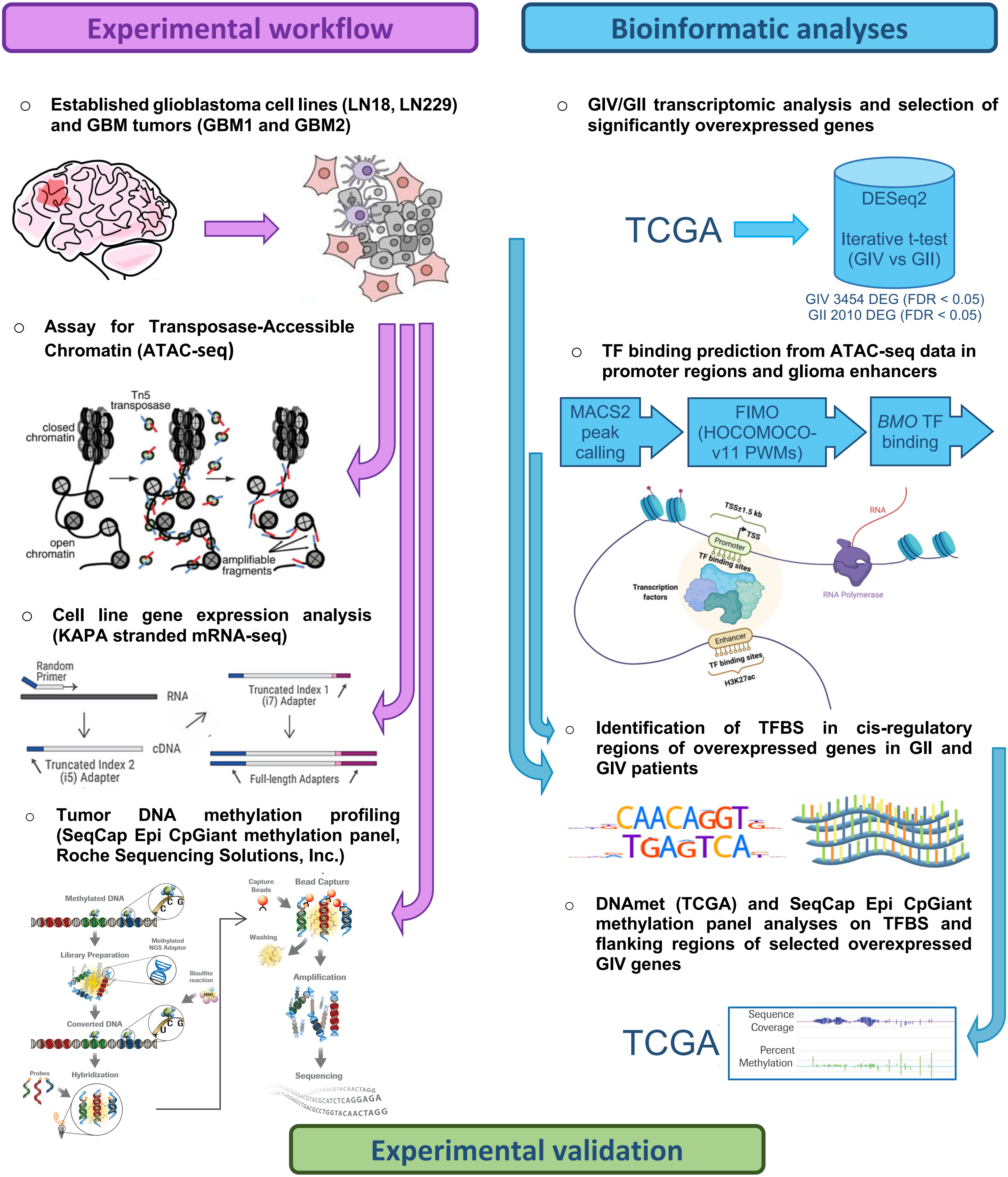
Experimental workflow
Bioinformatic analyses
Established glioblastoma cell lines (LN18, LN229) and GBM tumors (GBM1 and GBM2)
GIV/GII transcriptomic analysis and selection of significantly overexpressed genes
TCGA
DESeq2
Iterative t-test (GIV vs GII)
GIV 3454 DEG (FDR < 0.05)
GII 2010 DEG (FDR < 0.05)
Assay for Transposase-Accessible Chromatin (ATAC-seq)
Tn5 transposase
closed chromatin
open chromatin
amplifiable fragments
TF binding prediction from ATAC-seq data in promoter regions and glioma enhancers
MACS2 peak calling
FIMO (HOCOMOCO-v11 PWMs)
BMO TF binding
TSS±1.5 kb
TSS
Promoter
TF binding sites
Transcription factors
RNA
RNA Polymerase
Enhancer
TF binding sites
H3K27ac
Cell line gene expression analysis (KAPA stranded mRNA-seq)
Random Primer
RNA
cDNA
Truncated Index 1 (i7) Adapter
Truncated Index 2 (i5) Adapter
Full-length Adapters
Identification of TFBS in cis-regulatory regions of overexpressed genes in GII and GIV patients
CAACAGGT
TGAGTCA
Tumor DNA methylation profiling (SeqCap Epi CpGiant methylation panel, Roche Sequencing Solutions, Inc.)
Methylated DNA
Library Preparation
Methylated NGS Adaptor
Bisulfite reaction
Converted DNA
Probes
Hybridization
Capture Beads
Bead Capture
Washing
Amplification
Sequencing
DNAmet (TCGA) and SeqCap Epi CpGiant methylation panel analyses on TFBS and flanking regions of selected overexpressed GIV genes
TCGA
Sequence Coverage
Percent Methylation
Experimental validation

**Figure S1.** Study's experimental and bioinformatic effort depicted in a schematic diagram.

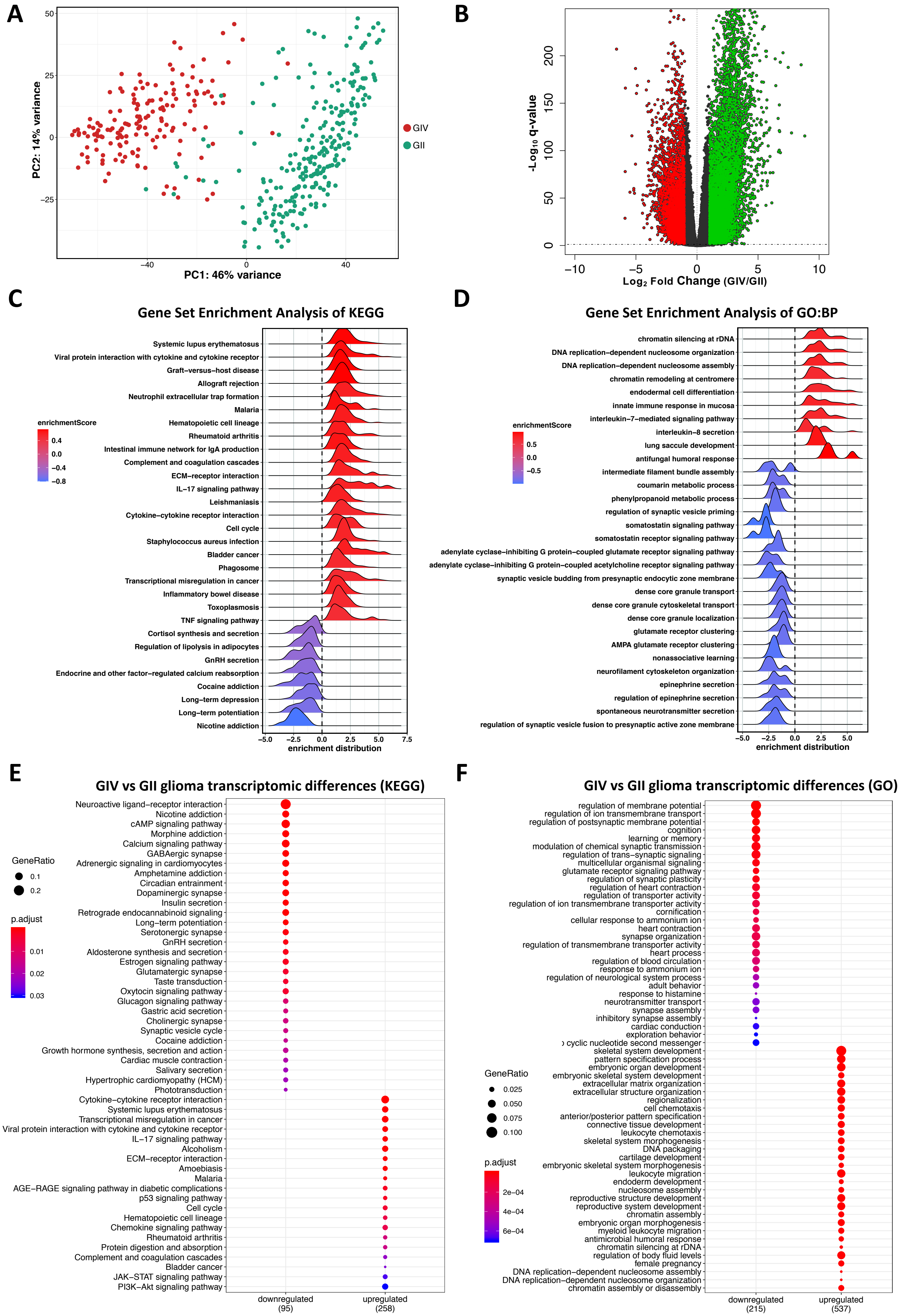

A
PC2: 14% variance
PC1: 46% variance
GIV
GII
B
-Log10 q-value
Log2 Fold Change (GIV/GII)
C
Gene Set Enrichment Analysis of KEGG
enrichmentScore
Systemic lupus erythematosus
Viral protein interaction with cytokine and cytokine receptor
Graft−versus−host disease
Allograft rejection
Neutrophil extracellular trap formation
Malaria
Hematopoietic cell lineage
Rheumatoid arthritis
Intestinal immune network for IgA production
Complement and coagulation cascades
ECM−receptor interaction
IL−17 signaling pathway
Leishmaniasis
Cytokine−cytokine receptor interaction
Cell cycle
Staphylococcus aureus infection
Bladder cancer
Phagosome
Transcriptional misregulation in cancer
Inflammatory bowel disease
Toxoplasmosis
TNF signaling pathway
Cortisol synthesis and secretion
Regulation of lipolysis in adipocytes
GnRH secretion
Endocrine and other factor−regulated calcium reabsorption
Cocaine addiction
Long−term depression
Long−term potentiation
Nicotine addiction
enrichment distribution
D
Gene Set Enrichment Analysis of GO:BP
enrichmentScore
chromatin silencing at rDNA
DNA replication−dependent nucleosome organization
DNA replication−dependent nucleosome assembly
chromatin remodeling at centromere
endodermal cell differentiation
innate immune response in mucosa
interleukin−7−mediated signaling pathway
interleukin−8 secretion
lung saccule development
antifungal humoral response
intermediate filament bundle assembly
coumarin metabolic process
phenylpropanoid metabolic process
regulation of synaptic vesicle priming
somatostatin signaling pathway
somatostatin receptor signaling pathway
adenylate cyclase−inhibiting G protein−coupled glutamate receptor signaling pathway
adenylate cyclase−inhibiting G protein−coupled acetylcholine receptor signaling pathway
synaptic vesicle budding from presynaptic endocytic zone membrane
dense core granule transport
dense core granule cytoskeletal transport
dense core granule localization
glutamate receptor clustering
AMPA glutamate receptor clustering
nonassociative learning
neurofilament cytoskeleton organization
epinephrine secretion
regulation of epinephrine secretion
spontaneous neurotransmitter secretion
regulation of synaptic vesicle fusion to presynaptic active zone membrane
enrichment distribution
E
GIV vs GII glioma transcriptomic differences (KEGG)
GeneRatio
p.adjust
Neuroactive ligand−receptor interaction
Nicotine addiction
cAMP signaling pathway
Morphine addiction
Calcium signaling pathway
GABAergic synapse
Adrenergic signaling in cardiomyocytes
Amphetamine addiction
Circadian entrainment
Dopaminergic synapse
Insulin secretion
Retrograde endocannabinoid signaling
Long−term potentiation
Serotonergic synapse
GnRH secretion
Aldosterone synthesis and secretion
Estrogen signaling pathway
Glutamatergic synapse
Taste transduction
Oxytocin signaling pathway
Glucagon signaling pathway
Gastric acid secretion
Cholinergic synapse
Synaptic vesicle cycle
Cocaine addiction
Growth hormone synthesis, secretion and action
Cardiac muscle contraction
Salivary secretion
Hypertrophic cardiomyopathy (HCM)
Phototransduction
Cytokine−cytokine receptor interaction
Systemic lupus erythematosus
Transcriptional misregulation in cancer
Viral protein interaction with cytokine and cytokine receptor
IL−17 signaling pathway
Alcoholism
ECM−receptor interaction
Amoebiasis
Malaria
AGE−RAGE signaling pathway in diabetic complications
p53 signaling pathway
Cell cycle
Hematopoietic cell lineage
Chemokine signaling pathway
Rheumatoid arthritis
Protein digestion and absorption
Complement and coagulation cascades
Bladder cancer
JAK−STAT signaling pathway
PI3K−Akt signaling pathway
downregulated (95)
upregulated (258)
F
GIV vs GII glioma transcriptomic differences (GO)
GeneRatio
p.adjust
regulation of membrane potential
regulation of ion transmembrane transport
regulation of postsynaptic membrane potential
cognition
learning or memory
modulation of chemical synaptic transmission
regulation of trans−synaptic signaling
multicellular organismal signaling
glutamate receptor signaling pathway
regulation of synaptic plasticity
regulation of heart contraction
regulation of transporter activity
regulation of ion transmembrane transporter activity
cornification
cellular response to ammonium ion
heart contraction
synapse organization
regulation of transmembrane transporter activity
heart process
regulation of blood circulation
response to ammonium ion
regulation of neurological system process
adult behavior
response to histamine
neurotransmitter transport
synapse assembly
inhibitory synapse assembly
cardiac conduction
exploration behavior
cyclic nucleotide second messenger
skeletal system development
pattern specification process
embryonic organ development
embryonic skeletal system development
extracellular matrix organization
extracellular structure organization
regionalization
cell chemotaxis
anterior/posterior pattern specification
connective tissue development
leukocyte chemotaxis
skeletal system morphogenesis
DNA packaging
cartilage development
embryonic skeletal system morphogenesis
leukocyte migration
endoderm development
nucleosome assembly
reproductive structure development
reproductive system development
chromatin assembly
embryonic organ morphogenesis
myeloid leukocyte migration
antimicrobial humoral response
chromatin silencing at rDNA
regulation of body fluid levels
female pregnancy
DNA replication−dependent nucleosome assembly
DNA replication−dependent nucleosome organization
chromatin assembly or disassembly
downregulated (215)
upregulated (537)

**Figure S2.** Transcriptomic analysis of high- and low-grade gliomas from the TCGA data (248 GII gliomas and 160 GIV gliomas). **(A)** Principal component analysis (PCA) was used to plot TCGA samples. The first two principal components (PCs) are plotted and colored according to the patient's glioma grade. The TCGA's normalized RNA-seq expression data were used to perform PCA. The axis label displays the percentage of variation accounted for by each principal component. **(B)** Volcano plot depicting the relevant gene expression differences between glioma grades (GIV vs GII). Green and red dots represent statistically significant up-regulated genes (DESeq2 methods, padj<0.05) in GIV gliomas or GII gliomas with logFC>1. The q-value threshold is indicated by a dotted horizontal line. **(C, D)** Gene set enrichment analysis (GSEA) of highly expressed genes in GIV glioma (red, 10% quantile categories) versus GII glioma (blue, 90% quantile categories) using the **(C)** KEGG and **(D)** Gene Ontology: Biological Processes databases. **(E, F)** Pathway enrichment analysis of differentially expressed genes (up logFC>2.5 and padj<0.01; down logFC<-2.5 and padj<0.01) reveals regulatory pathways in GIV glioma versus GII glioma. Raw counts were pre-filtered (> 5 reads within the cohort), and the Benjamini–Hochberg (BH) procedure was used to correct for multiple testing.

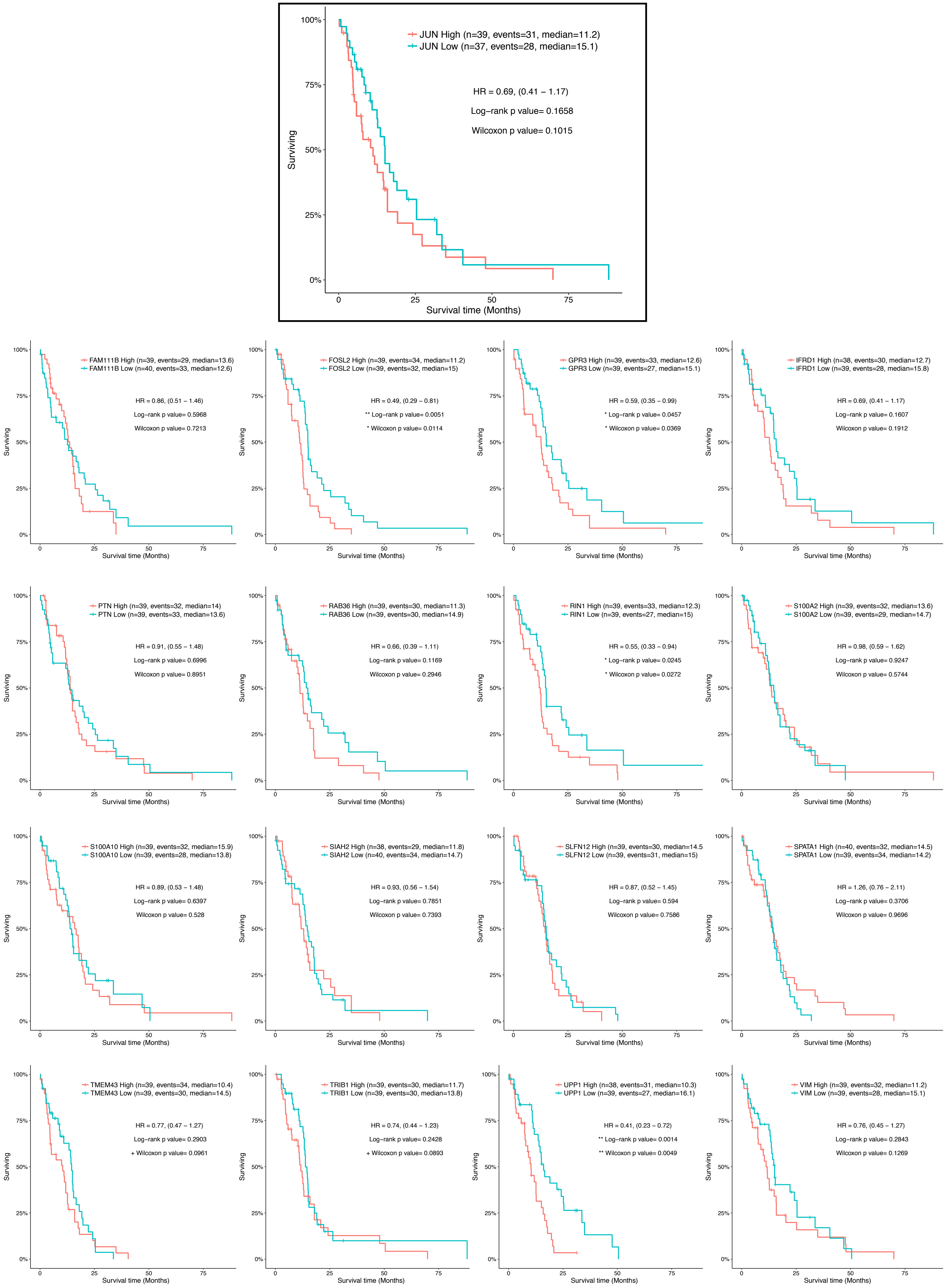
JUN High (n=39, events=31, median=11.2)
JUN Low (n=37, events=28, median=15.1)
HR = 0.69, (0.41 – 1.17)
Log−rank p value= 0.1658
Wilcoxon p value= 0.1015
Surviving
Survival time (Months)
FAM111B High (n=39, events=29, median=13.6)
FAM111B Low (n=40, events=33, median=12.6)
HR = 0.86, (0.51 – 1.46)
Log−rank p value= 0.5968
Wilcoxon p value= 0.7213
FOSL2 High (n=39, events=34, median=11.2)
FOSL2 Low (n=39, events=32, median=15)
HR = 0.49, (0.29 – 0.81)
** Log−rank p value= 0.0051
* Wilcoxon p value= 0.0114
GPR3 High (n=39, events=33, median=12.6)
GPR3 Low (n=39, events=27, median=15.1)
HR = 0.59, (0.35 – 0.99)
* Log−rank p value= 0.0457
* Wilcoxon p value= 0.0369
IFRD1 High (n=38, events=30, median=12.7)
IFRD1 Low (n=39, events=28, median=15.8)
HR = 0.69, (0.41 – 1.17)
Log−rank p value= 0.1607
Wilcoxon p value= 0.1912
PTN High (n=39, events=32, median=14)
PTN Low (n=39, events=33, median=13.6)
HR = 0.91, (0.55 – 1.48)
Log−rank p value= 0.6996
Wilcoxon p value= 0.8951
RAB36 High (n=39, events=30, median=11.3)
RAB36 Low (n=39, events=30, median=14.9)
HR = 0.66, (0.39 – 1.11)
Log−rank p value= 0.1169
Wilcoxon p value= 0.2946
RIN1 High (n=39, events=33, median=12.3)
RIN1 Low (n=39, events=27, median=15)
HR = 0.55, (0.33 – 0.94)
* Log−rank p value= 0.0245
* Wilcoxon p value= 0.0272
S100A2 High (n=39, events=32, median=13.6)
S100A2 Low (n=39, events=29, median=14.7)
HR = 0.98, (0.59 – 1.62)
Log−rank p value= 0.9247
Wilcoxon p value= 0.5744
S100A10 High (n=39, events=32, median=15.9)
S100A10 Low (n=39, events=28, median=13.8)
HR = 0.89, (0.53 – 1.48)
Log−rank p value= 0.6397
Wilcoxon p value= 0.528
SIAH2 High (n=38, events=29, median=11.8)
SIAH2 Low (n=40, events=34, median=14.7)
HR = 0.93, (0.56 – 1.54)
Log−rank p value= 0.7851
Wilcoxon p value= 0.7393
SLFN12 High (n=39, events=30, median=14.5
SLFN12 Low (n=39, events=31, median=15)
HR = 0.87, (0.52 – 1.45)
Log−rank p value= 0.594
Wilcoxon p value= 0.7586
SPATA1 High (n=40, events=32, median=14.5)
SPATA1 Low (n=39, events=34, median=14.2)
HR = 1.26, (0.76 – 2.11)
Log−rank p value= 0.3706
Wilcoxon p value= 0.9696
TMEM43 High (n=39, events=34, median=10.4)
TMEM43 Low (n=39, events=30, median=14.5)
HR = 0.77, (0.47 – 1.27)
Log−rank p value= 0.2903
+ Wilcoxon p value= 0.0961
TRIB1 High (n=39, events=30, median=11.7)
TRIB1 Low (n=39, events=30, median=13.8)
HR = 0.74, (0.44 – 1.23)
Log−rank p value= 0.2428
+ Wilcoxon p value= 0.0893
UPP1 High (n=38, events=31, median=10.3)
UPP1 Low (n=39, events=27, median=16.1)
HR = 0.41, (0.23 – 0.72)
** Log−rank p value= 0.0014
** Wilcoxon p value= 0.0049
VIM High (n=39, events=32, median=11.2)
VIM Low (n=39, events=28, median=15.1)
HR = 0.76, (0.45 – 1.27)
Log−rank p value= 0.2843
Wilcoxon p value= 0.1269

**Figure S3.** Glioblastomas (GBM-TCGA) median overall survival (OS) estimation based on gene expression for c-Jun target genes using the Kaplan-Meier method (cut-off defined by splitting datasets in 25% lower expressing vs 25% higher expressing patients). Vertical marks indicate censorship, the number of patients and median survival in months per group are shown, and the hazard ratio (HR) is defined. The log-rank test and Wilcoxon p-values are displayed (*p-value < 0.05, **p-value < 0.01 and ***p-value < 0.001).

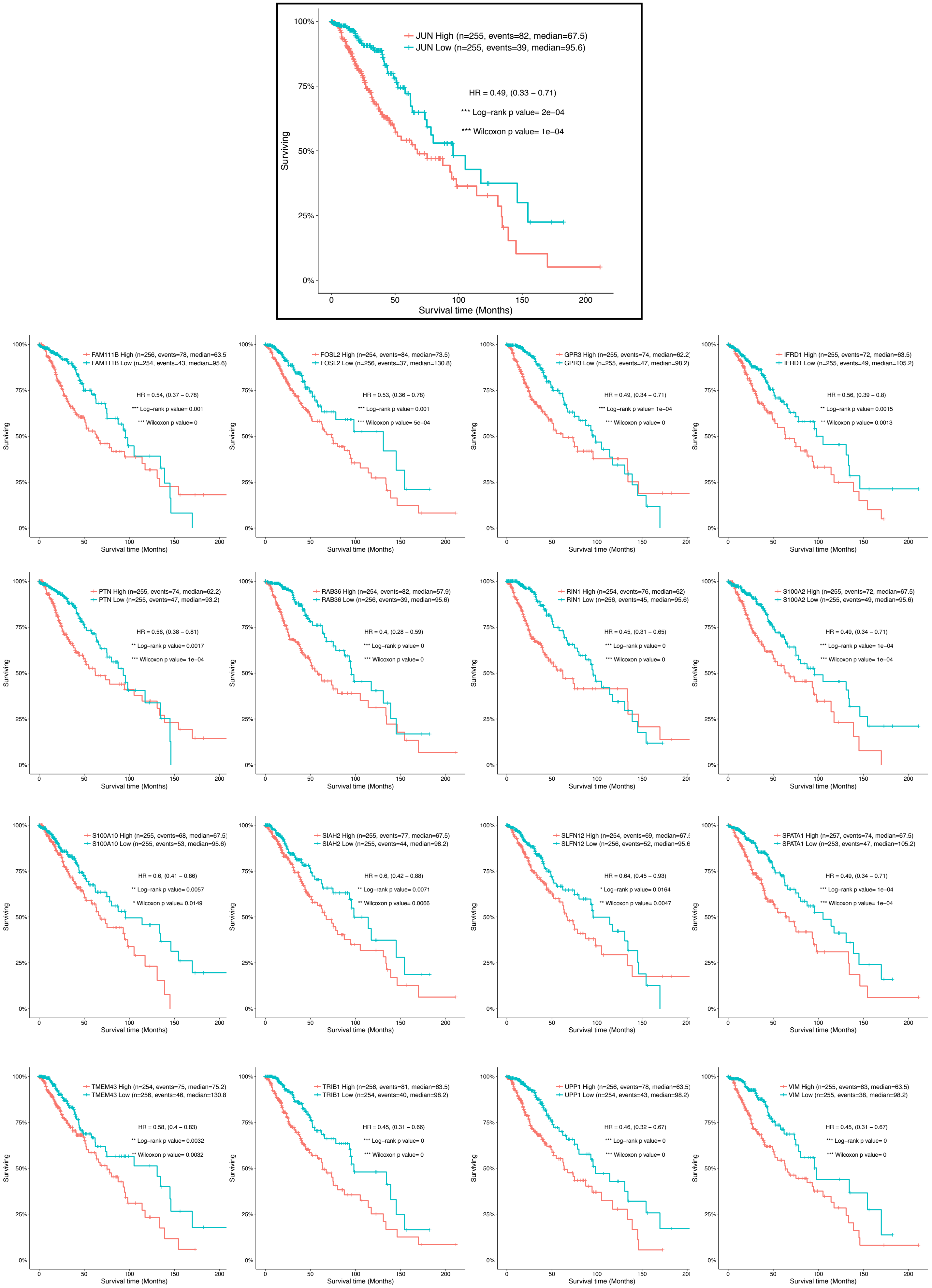

JUN High (n=255, events=82, median=67.5)
JUN Low (n=255, events=39, median=95.6)
HR = 0.49, (0.33 − 0.71)
*** Log−rank p value= 2e−04
*** Wilcoxon p value= 1e−04
Surviving
Survival time (Months)
FAM111B High (n=256, events=78, median=63.5
FAM111B Low (n=254, events=43, median=95.6)
HR = 0.54, (0.37 − 0.78)
*** Log−rank p value= 0.001
*** Wilcoxon p value= 0
FOSL2 High (n=254, events=84, median=73.5)
FOSL2 Low (n=256, events=37, median=130.8)
HR = 0.53, (0.36 − 0.78)
*** Log−rank p value= 0.001
*** Wilcoxon p value= 5e−04
GPR3 High (n=255, events=74, median=62.2
GPR3 Low (n=255, events=47, median=98.2)
HR = 0.49, (0.34 − 0.71)
*** Log−rank p value= 1e−04
*** Wilcoxon p value= 0
IFRD1 High (n=255, events=72, median=63.5)
IFRD1 Low (n=255, events=49, median=105.2)
HR = 0.56, (0.39 − 0.8)
** Log−rank p value= 0.0015
** Wilcoxon p value= 0.0013
PTN High (n=255, events=74, median=62.2)
PTN Low (n=255, events=47, median=93.2)
HR = 0.56, (0.38 − 0.81)
** Log−rank p value= 0.0017
*** Wilcoxon p value= 1e−04
RAB36 High (n=254, events=82, median=57.9)
RAB36 Low (n=256, events=39, median=95.6)
HR = 0.4, (0.28 − 0.59)
*** Log−rank p value= 0
*** Wilcoxon p value= 0
RIN1 High (n=254, events=76, median=62)
RIN1 Low (n=256, events=45, median=95.6)
HR = 0.45, (0.31 − 0.65)
*** Log−rank p value= 0
*** Wilcoxon p value= 0
S100A2 High (n=255, events=72, median=67.5)
S100A2 Low (n=255, events=49, median=95.6)
HR = 0.49, (0.34 − 0.71)
*** Log−rank p value= 1e−04
*** Wilcoxon p value= 1e−04
S100A10 High (n=255, events=68, median=67.5)
S100A10 Low (n=255, events=53, median=95.6)
HR = 0.6, (0.41 − 0.86)
** Log−rank p value= 0.0057
* Wilcoxon p value= 0.0149
SIAH2 High (n=255, events=77, median=67.5)
SIAH2 Low (n=255, events=44, median=98.2)
HR = 0.6, (0.42 − 0.88)
** Log−rank p value= 0.0071
** Wilcoxon p value= 0.0066
SLFN12 High (n=254, events=69, median=67.5
SLFN12 Low (n=256, events=52, median=95.6
HR = 0.64, (0.45 − 0.93)
* Log−rank p value= 0.0164
** Wilcoxon p value= 0.0047
SPATA1 High (n=257, events=74, median=67.5)
SPATA1 Low (n=253, events=47, median=105.2)
HR = 0.49, (0.34 − 0.71)
*** Log−rank p value= 1e−04
*** Wilcoxon p value= 1e−04
TMEM43 High (n=254, events=75, median=75.2)
TMEM43 Low (n=256, events=46, median=130.8
HR = 0.58, (0.4 − 0.83)
** Log−rank p value= 0.0032
** Wilcoxon p value= 0.0032
TRIB1 High (n=256, events=81, median=63.5)
TRIB1 Low (n=254, events=40, median=98.2)
HR = 0.45, (0.31 − 0.66)
*** Log−rank p value= 0
*** Wilcoxon p value= 0
UPP1 High (n=256, events=78, median=63.5)
UPP1 Low (n=254, events=43, median=98.2)
HR = 0.46, (0.32 − 0.67)
*** Log−rank p value= 0
*** Wilcoxon p value= 0
VIM High (n=255, events=83, median=63.5)
VIM Low (n=255, events=38, median=98.2)
HR = 0.45, (0.31 − 0.67)
*** Log−rank p value= 0
*** Wilcoxon p value= 0

**Figure S4.** Low-Grade Gliomas (LGG-TCGA) median overall survival (OS) estimation based on gene expression for c-Jun target genes using the Kaplan-Meier method (cut-off defined by splitting datasets in 25% lower expressing vs 25% higher expressing patients). Vertical marks indicate censorship, the number of patients and median survival in months per group are shown, and the hazard ratio (HR) is defined. The log-rank test and Wilcoxon p-values are displayed (*p-value < 0.05, **p-value < 0.01 and ***p-value < 0.001).

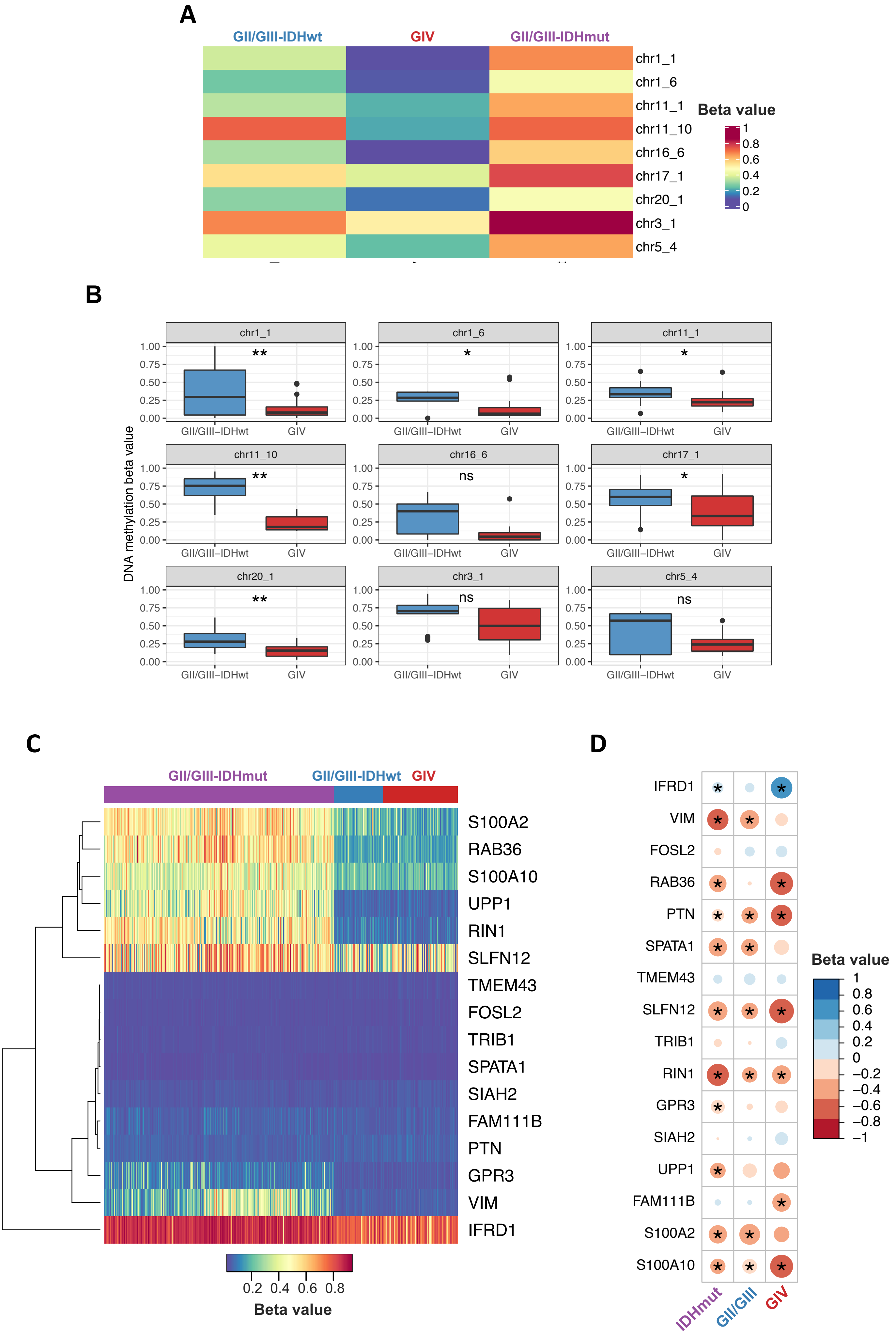
A
GII/GIII-IDHwt
GIV
GII/GIII-IDHmut
chr1_1
chr1_6
chr11_1
chr11_10
chr16_6
chr17_1
chr20_1
chr3_1
chr5_4
Beta value
1
0.8
0.6
0.4
0.2
0
B
DNA methylation beta value
chr1_1
**
chr1_6
*
chr11_1
*
chr11_10
**
chr16_6
ns
chr17_1
*
chr20_1
**
chr3_1
ns
chr5_4
ns
1.00
0.75
0.50
0.25
0.00
GII/GIII–IDHwt
GIV
C
GII/GIII-IDHmut
GII/GIII-IDHwt
GIV
S100A2
RAB36
S100A10
UPP1
RIN1
SLFN12
TMEM43
FOSL2
TRIB1
SPATA1
SIAH2
FAM111B
PTN
GPR3
VIM
IFRD1
0.2
0.4
0.6
0.8
Beta value
D
IFRD1
VIM
FOSL2
RAB36
PTN
SPATA1
TMEM43
SLFN12
TRIB1
RIN1
GPR3
SIAH2
UPP1
FAM111B
S100A2
S100A10
IDHmut
GII/GIII
GIV
Beta value
1
0.8
0.6
0.4
0.2
0
−0.2
−0.4
−0.6
−0.8
−1

**Figure S5**. DNA methylation pattern of c-Jun regulated genes in human gliomas. **(A)** Heatmap represents regional DNA methylation differences (C-rich regions overlapping c-Jun motifs) from putative glioma enhancers. **(B)** Boxplots depict C-rich regions overlapping c-Jun motifs having significantly different DNA methylation pattern among glioma groups (*FDR < 0.05, **FDR < 0.01 and ***FDR < 0.001). **(C)** DNA methylation pattern in the promoters (median beta value from the +2 kb/-500 bp relative to TSS) of joined LGG and GBM TCGA datasets (GII, III and IV of WHO 2016 classification). **(D)** Correlation plot of DNA methylation from the promoters (as in panel C) correlated with gene expression. Significant (FDR<0.05) correlations were marked with an asterisk.

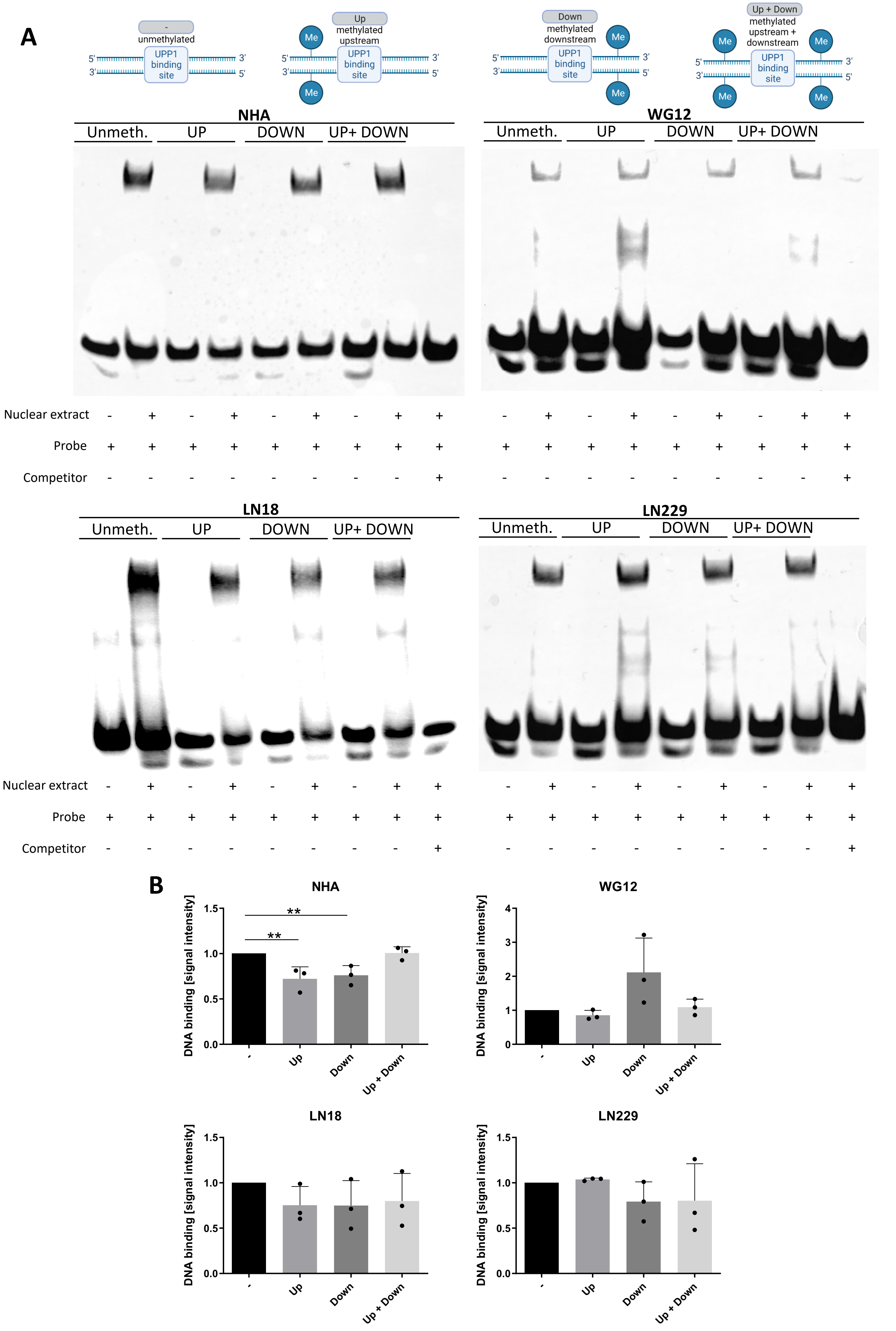

A
- unmethylated
UPP1 binding site
Up methylated upstream
Down methylated downstream
Up + Down methylated upstream + downstream
Me
5'
3'
NHA
WG12
LN18
LN229
Unmeth.
UP
DOWN
UP+ DOWN
Nuclear extract
Probe
Competitor
B
DNA binding [signal intensity]
**
-
Up
Down
Up + Down

**Figure S6.** EMSA for nuclear extracts employing probes with different methylation patterns. **(A)** Each EMSA compares binding of nuclear protein extract to DNA of probes containing the genomic sequence matching c-Jun motif (ACTCAGTGAA) within the *UPP1* promoter in different cell lines. Nuclear protein and DNA complex formation was compared for 4 different primers: unmethylated, upstream methylated, downstream methylated and up- and down-stream methylated. **(B)** Data were normalized to unmethylated oligonucleotides and p-values were calculated using the one-way ANOVA with Dunnett's post-hoc test, $^{**}p < 0.01$, ±SD.

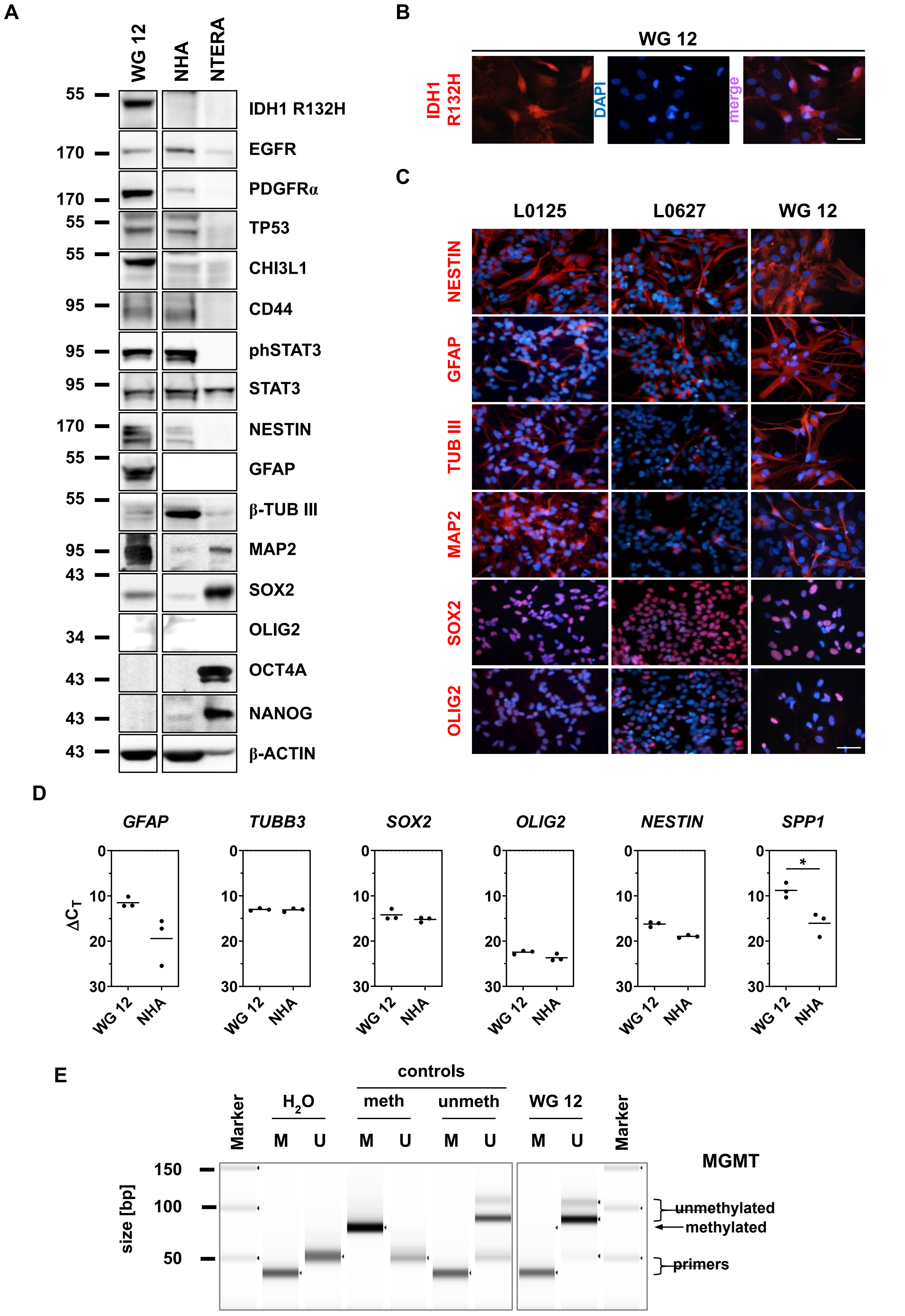
A
WG 12
NHA
NTERA
55
170
170
55
55
95
95
95
170
55
55
95
43
34
43
43
43
IDH1 R132H
EGFR
PDGFRα
TP53
CHI3L1
CD44
phSTAT3
STAT3
NESTIN
GFAP
β-TUB III
MAP2
SOX2
OLIG2
OCT4A
NANOG
β-ACTIN
B
WG 12
IDH1 R132H
DAPI
merge
C
L0125
L0627
WG 12
NESTIN
GFAP
TUB III
MAP2
SOX2
OLIG2
D
GFAP
TUBB3
SOX2
OLIG2
NESTIN
SPP1
ΔC_T
0
10
20
30
*
WG 12
NHA
E
controls
Marker
H2O
meth
unmeth
WG 12
Marker
M
U
size [bp]
150
100
50
MGMT
unmethylated
methylated
primers

**Figure S7.** Characterization of patient-derived primary WG12 cell line. **(A)** Western Blot analysis of pluripotency (OCT4A, NANOG), neural stem/progenitor (NESTIN, SOX2, OLIG2), astrocytic (GFAP) and neuronal (MAP2, b-Tubulin III) markers in WG12 cells. Other proteins represent the markers of glioblastoma subtypes: CHI3L1, phSTAT3, CD44 (mesenchymal); EGFR (classical); PDGFRα, TP53, IDH1 R132H, (proneural) according to Verhaak classification. NHA cells serve as a non-malignant control, whereas NTERA cells as control with stemness phenotype. β-actin and STAT3 were used as a loading control. (**B)** Immunofluorescent staining of IDH1 R132H mutation in WG12 cells. **(C)** Immunostaining of neural stem/progenitor (NESTIN, OLIG2, SOX2) and differentiation (GFAP, MAP2, β-Tubulin III) markers in WG12 cells. Serum-differentiated L0125 and L0627 cells serve as a control for stemness and differentiation properties. **(D)** Expression of neural stem/progenitor (*SOX2*, *OLIG2*, *NESTIN*, *SPP1*) and differentiation (*GFAP*, *TUBB3*) markers in WG12 and in control NHA cells. **E** *MGMT* promoter methylation status analysis in glioma primary WG12 cells.
